# Causal contributions of neural noise to left frontal networks subtending the modulation of conscious visual perception in the human brain

**DOI:** 10.1101/2021.11.22.469553

**Authors:** Chloé Stengel, Julià L. Amengual, Tristan Moreau, Antoni Valero-Cabré

## Abstract

For several decades, the field of human neurophysiology has focused on the role played by cortical oscillations in enabling brain function underpinning behaviors. In parallel, a less visible but robust body of work on the stochastic resonance phenomenon has also theorized contributions of neural noise − hence more heterogeneous, complex and less predictable activity − in brain coding. The latter notion has received indirect causal support via improvements of visual function during non-regular or random brain stimulation patterns. Nonetheless, direct evidence demonstrating an impact of brain stimulation on direct measures of neural noise is still lacking. Here we evaluated the impact of three *non frequency-specific* TMS bursts, compared to a control pure high-beta TMS rhythm, delivered to the left FEF during a visual detection task, on the heterogeneity, predictability and complexity of ongoing brain activity recorded with scalp EEG. Our data showed surprisingly that the three *non frequency-specific* TMS patterns did not prevent a build-up of local high-beta activity. Nonetheless, they increased power across broader or in multiple frequency bands compared to control purely *rhythmic* high-beta bursts tested along. Importantly, *non frequency-specific* patterns enhanced signal entropy over multiple time-scales, suggesting higher complexity and an overall induction of higher levels of cortical noise than *rhythmic* TMS bursts. Our outcomes provide indirect evidence on a potential modulatory role played by sources of stochastic noise on brain oscillations and synchronization. Additionally, they pave the way towards the development of novel neurostimulation approaches to manipulate cortical sources of noise and further investigate their causal role in neural coding.

## Introduction

Uncovering the neural basis of cognition demands a detailed characterization in time and space of the neural patterns encoding specific brain states and cognitive functions, as well as plausible mechanistic models supporting such associations. Traditionally, research effort has focused on characterizing the role of highly predictable and low complexity neural signals, known as brain oscillations, and has devoted efforts to exploring the coding role of some of their features such as frequency, phase or local and interregional coherence measures.

In the domain of visuospatial attention, correlational studies in animal models (Buschman & Miller, 2007; Saalmann et al., 2007) and healthy humans (Gross et al., 2004; Hipp et al., 2011; Phillips & Takeda, 2009; Rodriguez et al., 1999) have associated fronto-parietal high-beta oscillations with a top-down allocation of spatial attention and the modulation of visual perception. Causal confirmation of such observations has come from rhythmic Transcranial Magnetic Stimulation (TMS) studies in which the entrainment of high-beta frequencies on the right Frontal Eye Field (FEF), a node of the bilaterally distributed dorsal attention network (Corbetta & Shulman, 2002), and associated parietal systems enhanced visual sensitivity (Chanes et al., 2013; Quentin et al., 2015; Stengel et al., 2021; Vernet et al., 2019).

Nonetheless, brain oscillations are not the only recordable neural signals with a potential to contribute to the top-down modulation of visual perception for incoming visual stimuli. Indeed, non frequency-specific, broader-band, less predictable and more complex activity, proven ubiquitous in EEG recordings and generally referred to as neural ‘noise’, has garnered increased attention, particularly with regards to its ability to enable cognitive processes underpinning human behavior. Particularly relevant for the field of visual perception, the framework posed by Stochastic Resonance (SR) theorized several decades ago that optimal levels of stochastic noise added to sub-threshold signals generated by non-linear systems may boost the saliency and detectability of weak stimuli (see Moss et al., 2004 for a review). Such initial hypotheses were substantiated by experimental evidence showing the benefits of externally added stochastic noise in signal detection by peripheral receptors (Collins et al., 1996; Cordo et al., 1996; Douglass et al., 1993) or its influence on the processing of neural signals by cortical systems (Groen & Wenderoth, 2016; Iliopoulos et al., 2014; Kitajo et al., 2003; Lugo et al., 2008; Manjarrez et al., 2007).

Regarding coding strategies subtending top-down modulatory mechanisms for conscious visual detection, causal evidence has suggested dissimilarity in neural coding used by frontal areas of the right and left hemispheres. More specifically, at difference with findings in the right Frontal Eye Fields (FEF) showing increases of visual sensitivity with the entrainment of rhythmic 30 Hz activity, *non frequency-specific* TMS patterns delivered prior to target onset to its left counterpart resulted in unexpected visual perception improvements (Chanes et al., 2015). Inspired by the Stochastic Resonance framework, these results ignited speculation that *non frequency-specific* TMS bursts may enhance conscious perception by increasing levels of neural ‘noise’ in ongoing left frontal activity. However, electrophysiological analyses focused on non-predictable neural activity, instead of rhythmic oscillatory fluctuations, were required to verify such hypothesis and, on such basis, build an accurate interpretation of the causal role played by sources of internal ‘noise’ on brain function, including mechanisms enabling the top-down modulation of perception in attention orienting networks.

We here applied concurrent TMS-EEG recordings and tested if, compared to pure high-beta 30 Hz *rhythmic* activity, three different types of *non frequency-specific* TMS bursts (*non-uniform rhythmic, random and irregular*) all delivered pre-target onset during performance of a lateralized visual detection task on near-threshold targets, would differentially modulate patterns of neural activity occurring in left fronto-parietal nodes of the dorsal visuo-spatial attention network. Three specific predictions substantiated by specific planned analyses were made. First, we hypothesized that *non frequency-specific* TMS patterns would increase the power of oscillations in a broader range of frequencies than *rhythmic* high-beta bursts. Second, we predicted that the former broadband effects would also manifest as increases in the level of non-predictable activity, hence analyzed via measures of signal entropy and complexity extracted from the recorded EEG signals. Third and last, we anticipated that well-dosed levels of neural noise induced by some but not all of the three *non frequency-specific* TMS (and certainly not with *rhythmic* TMS) would potentially drive improvements of visual perception in healthy participants.

## Materials and Methods

### Participants

A group of 15 right-handed participants (9 women and 6 males) aged between 21 and 45 years old (29 ± 6, mean ± SD) took part in the current study. Participants reported no history of neurological disorders and had normal or corrected-to-normal vision. All of them voluntarily consented to participate in the study and signed a consent form. The research protocol including all the interventions of this study (C08-45/C14-17) was sponsored by the INSERM (Institut National de la Santé et la Recherche Médicale) and approved by an Institutional Review Board, the Comité de Protection des Personnes (CPP), Ile de France V.

### Conscious visual detection paradigm

Similar tasks have been employed in prior publications by our research group (see Chanes et al., 2013, 2015; Quentin et al., 2015; Vernet et al., 2019). The presentation of visual stimuli was controlled by an in-house MATLAB 2012b (Mathworks) script using the Psychotoolbox extensions (Brainard, 1997) and synchronized with the delivery of the TMS pulses (see Fig. 1A for a schematic representation of the sequence of events during a trial). Participants were seated with their eye’s canthi positioned 57 cm away from the center of a computer screen. Trials started with a fixation screen that displayed a central fixation cross (size 0.5×0.5°) and a right and left rectangular placeholders (6.0 x 5.5°, drawn 8.5° away from the center of the screen) indicating the potential location of a visual target later in the trial. After an interval randomly jittered between 1000 and 1500 ms, the fixation cross became slightly larger (size 0.7×0.7°) during 66 ms to alert participants that the target would soon appear on the screen. Following an inter-stimulus-interval of 233 ms, a vertical low-contrast Gabor stimulus (0.5°/cycle sinusoidal spatial frequency, 0.6° exponential standard deviation) could appear for 33 ms in the center of one of the two placeholders with equal probability (40% of trials with left target, 40% of trials with right target, 20% catch trials with no target). During a calibration block prior to the beginning of the experimental session, Gabor contrast was adjusted for each participant to reach 50% detection rates.

**Figure 1.**
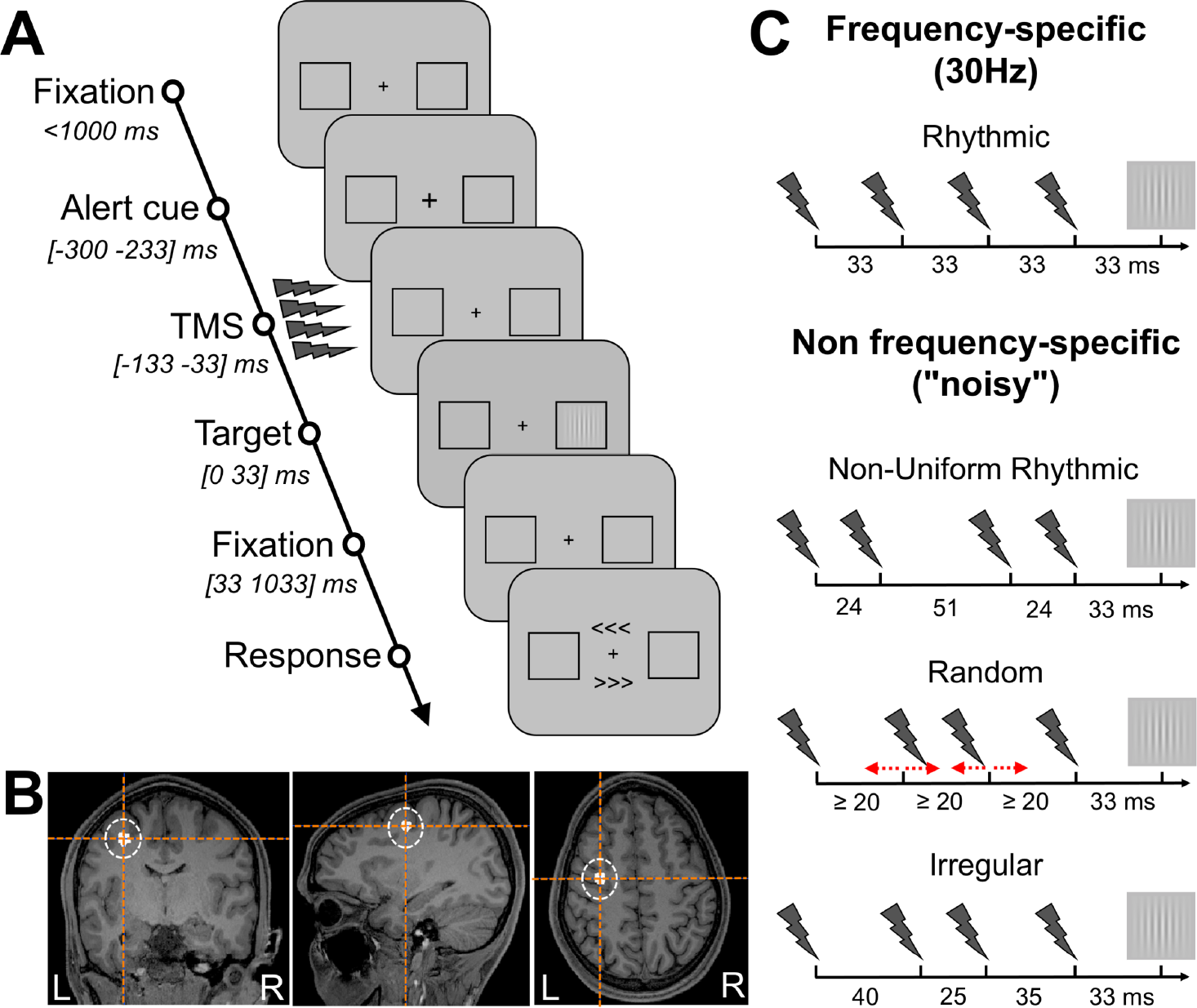
Visual detection task, targeted cortical region and TMS patterns. **(A)** Visual detection task performed by participants. After a period of fixation, a central cross became slightly larger to alert participants of an upcoming event. Then active or sham patterns of 30 Hz *rhythmic* or three *non frequency-specific* TMS were delivered to the left FEF prior to the presentation of a visual target (a near threshold 50% visibility Gabor) that could appear for a brief period of time (33 ms) at the center of a right or left placeholder. Participants were requested to indicate whether they did perceive a target or not (no/yes), and, when they consciously reported to have seen it, indicate where it appeared (right/left). Notice that in 20% of the trials (catch trials), no target was presented in any of the placeholders. (**B)** Coronal, axial and sagittal MRI sections from the frameless stereotaxic neuronavigation system showing the localization of the targeted left FEF (Talairach coordinates X=-32, Y=-2, Z=46) on a T1-3D MRI of a representative participant. **(C)** Schematic representation of the TMS patterns employed for active and sham stimulation. 30 Hz *rhythmic* pattern (designed to entrain oscillatory activity at the input frequency) and the three *non frequency-specific* TMS patterns (*non-uniform rhythmic*, *random* and *irregular* 4-pulse bursts) designed to locally generate different levels of neural noise in the left FEF.

Participants were presented with a response screen 1000 ms after the Gabor target offset. They were asked to perform a detection task in which they had to report whether they saw a target and, in the case of an affirmative response, where had the target appeared (left/right of the fixation cross). The response screen consisted in two arrow-like signs (“>>>” and “<<<”) displayed above and below the central fixation cross. Participants were asked to report which arrow pointed towards the placeholder (right or left) in which they saw the target. The location of the arrows (above or below the fixation cross) was randomized across trials, preventing participants from preparing their motor response prior to the onset of the response window. Participants responded with the index, middle and thumb fingers of their left hand by pressing the ‘d’ letter key to select the upper arrow, ‘c’ letter key for the lower arrow or the space bar to signal that they had not seen any target. The response of the participant ended the trial.

The contrast of the visual target was adjusted to reach the individual threshold contrast for which each participant showed consistent 50% detection performance following a one-up/one-down staircase procedure. Gabor contrast and contrast steps were initially set at a level of 1 Michelson units of contrast and upon each reversal of response the contrast step was divided by two. Note that, regardless, the contrast of the target throughout the titration procedure was always kept between 1 and 0.005 Michelson units of contrast. A consistent estimation of the 50% conscious detection threshold contrast was reached when, after five consecutive trials, target contrast varied by less than 0.01 Michelson contrast units. The threshold was measured twice using this exact same procedure. If the two contrast thresholds differed by less than 0.01 Michelson contrast unit, the calibration block was terminated and the contrast used for the rest of the experimental session was the average between the two thresholds. If this criterion was not fulfilled, the threshold was determined again until two consecutive titrations yielded contrasts that varied by less than 0.01 Michelson contrast units. During the calibration block, participants received only sham TMS (see below for details on the TMS procedure).

Following the titration of Gabor contrast, participants underwent a training block in which they were given the opportunity to become familiar with stimulation by getting exposed to active TMS trials. The order of trials (leftward target, rightward target or catch trial with no target) and the stimulation condition (sham and active TMS, see below for details on the TMS procedure) was randomized for each sub-block of 20 trials. During the training block, at the end of each sub-block participants received feedback about some aspects of their performance, including the percentage of trials in which visual fixation was broken (see below for details on the procedure for control of centrale gaze fixation) and the percentage of incorrectly reported target positions. Participants were also alerted if their false alarm rate was higher than 50 %. The training block was terminated by the experimenter on the basis of individual performance. Experimental blocks were identical to training blocks except that participants received feedback only every two sub-blocks. Following the presentation of feedback on the screen, participants were allowed to take a short break (∼2 minutes). Experimental blocks consisted of 7 sub-blocks (140 trials total) and lasted approximately 20 minutes each.

### Behavioral data analyses

The trials of the detection task were classified into different categories according to the different types of participant’s responses. Trials in which a target was present and participants correctly reported its presence and location were classified as a ‘Hit’, whereas when such a target was reported as ‘unseen’ they were classified as a ‘Miss’. Catch trials in which no target was displayed were counted either as a ‘False Alarm’ if participants reported the presence of a target or as a ‘Correct Rejection’ if this was reported as ‘unseen’. Very rarely, participants correctly reported the presence of a target but signaled an incorrect (right or left) location for it. These trials were classified as ‘Errors’ and excluded from further analyses as it became impossible to determine if participants had seen a target in a location where no target had been presented (akin to a ‘False Alarm’ trial) or simply pressed the wrong key when asked to report the target location.

Following the Signal Detection Theory (SDT) approach, on the basis of the rate of ‘Hit’ (H) trials and ‘False Alarm’ (FA) trials, we extracted separate measures of target perception and late-stage decision-making processes when delivering a response (Green & Swets, 1966; Stanislaw & Todorov, 1999). First, we computed perceptual sensitivity (d’), which is a bias free measure of a participant’s ability to distinguish the presence of a target from noise. Second, we calculated the decision criterion (c) and the likelihood ratio (β) which are both measures of the response bias of participants. Indeed, in case of doubt, participants might be biased to be more prone to indicate they saw a target (liberal decision making) or, on the contrary, to be more likely to respond they did not see any target (conservative decision making) independently of how well they perceived it. These outcome measures were calculated as follows:

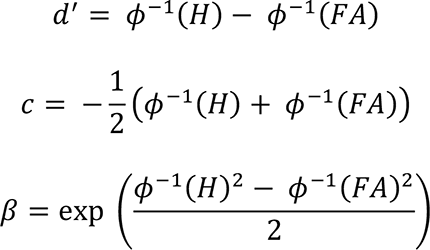

Where 𝜙^&’^ is the inverse of the normal cumulative distribution function. To avoid infinite values, a null rate of false alarms was corrected to 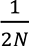 and a rate of hit trials of 1 was corrected to 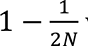 where N was the total number of trials on which each rate was calculated, following a well-established procedure (Macmillan & Creelman, 2004).

On the estimated values of d’, c and β, we performed a 2×2×4 repeated measure ANOVA with factors *Visual Field* (left, right visual target), *TMS Condition* (active, sham) and *TMS Pattern* (*rhythmic*, *non uniform rhythmic*, *random*, *irregular*, see below for details on TMS patterns).

### Control of central gaze fixation

In order to ensure reliable central fixation during the visual detection task, the position of both eyes was monitored with a remote eye-tracking system (Eyelink 1000, SR Research, sampling rate 1000 Hz) throughout the experimental session. If at any point between the onset of the alerting cue and the target offset, participants’ gaze was more than 2° away from the center of the fixation cross, the trial was aborted, labeled as incorrectly fixated and any data associated with it was excluded from analyses. In such cases, participants were alerted that they had violated fixation requirements with a message on the screen and the trial was re-randomized within the remaining trials to complete the sub-block and re-tested ulteriorly.

### Transcranial Magnetic Stimulation and neuronavigation procedures

Transcranial stimulation was delivered with a biphasic repetitive stimulator (SuperRapid^2^, Magstim) and a standard 70 mm diameter figure-of-eight TMS coil held tangentially on the skull over a region overlying the left Frontal Eye Field (FEF). TMS pulses were triggered via a high temporal resolution multichannel synchronization device (Master 8, AMPI, temporal resolution of 1 μsec) using TTL pulses. The position of the coil was tracked throughout the experiment with a neuronavigation system (Brainsight, Rogue Research). To this end, the left FEF was localized on individual T1-weighted MRI scans (3T Siemens MPRAGE, flip angle=9, TR=2300 ms, TE=4.18 ms, slice thickness=1mm) and labelled as a 5 mm radius spherical region of interest centered on Talairach coordinates x=-32, y=-2, z=46 (Paus, 1996) (Fig. 1B). The TMS coil was angled tangentially to the skull and held in a position ensuring the shortest Euclidian distance between the center of the TMS coil and the cortical region of interest. The coil handle was oriented ∼parallel to the central sulcus, at a ∼45° angle in a rostral to caudal and lateral to medial direction. Accurate coil positioning was monitored at all times during the experiment by an individual T1-3D MRI-based neuronavigation system which allowed investigators to keep the TMS coil within a ±3 mm radius from the center of the targeted cortical site, with identical angulation and tilting, throughout blocks and experimental sessions.

Sham stimulation was delivered through a round audio speaker (Mobi Wavemaster) attached to the upper flat surface of the TMS coil. During the simulation of a TMS pulse (i.e., the delivery of a sham TMS pulse), the speaker played a recording with the same acoustic properties as the characteristic clicking noise heard concomitantly with the delivery of an active magnetic pulse through a TMS coil. The audio file of a TMS pulse was generated by averaging the individual sound waveform of 100 single individual TMS pulse recordings (Aiwa CM-S32 stereo microphone). The envelope of the average waveform was then adjusted to emphasize high amplitude spikes at the beginning of the pulse so that, once replayed through our speaker, the sound became indistinguishable from the loud click produced by the TMS coil. The volume of the speaker was also adjusted to reproduce the same volume as an active TMS pulse. The precise onset timing of sham auditory pulses was handled by an in-house MATLAB script using the Psychotoolbox library extensions (Brainard, 1997), which was in control of the presentation of the events of the visual detection paradigm performed along.

Four different TMS patterns (Fig. 1C), all comprised of 4 TMS pulses spanning a window of 100 ms (between the onset of the 1^st^ pulse and the 4^th^ pulse) were delivered in our experiment: a *rhythmic* pattern with pulses regularly spaced in time to be delivered at a frequency of 30 Hz (i.e. with a fixed inter-pulse-interval of ∼33 ms) and designed to entrain high-beta cortical oscillations (Stengel et al., 2021; Vernet et al., 2019), and 3 *non frequency-specific* patterns, referred to as *non-uniform rhythmic*, *random* and *irregular* patterns, tailored to inject different levels of neural noise to neuronal assemblies hosted in the left FEF cortical region.

In the *non frequency-specific* TMS patterns, the timing of the 1^st^ and 4^th^ (hence last) pulse of each burst were kept identical to those of the *rhythmic* pattern. For the remaining patterns, the onset time of the two middle pulses (2^nd^ and 3^rd^ pulses) was shifted with regards to onset times in the *rhythmic* 30 Hz pattern to generate unequal inter-pulse intervals. In the *non-uniform rhythmic* pattern, the two middle pulses where anticipated and delayed by 9 ms, respectively. In the *random* pattern, the onset time of the 2nd and 3rd pulse was pseudo-randomly jittered to fall before or after their regular onset times in pure 30 Hz *rhythmic* patterns. Such jittering was constrained in two ways: first, in order to provide enough time for the TMS machine capacitors to recharge in-between pulses at the fixed TMS intensity employed in our experiments (see below for further details) a minimal inter-pulse interval of 20 ms had to be respected. Second, to ensure *random* patterns would never produce a perfectly regular 30 Hz frequency, the two middle pulses of each 4 pulse-burst were shifted at least 3 ms away from their onset time in the 30 Hz *rhythmic* pattern. Lastly, in *irregular* TMS patterns, the onset time of the two middle pulses (2^nd^ and 3^rd^ pulses) was set randomly within a 100 ms total burst duration, and respected the same limitations imposed to *random* patterns plus an additional constraint: the 3 inter-pulse intervals had to all have different durations. The onset times of the two middle pulses in *irregular* patterns were fixed for all trials. The *rhythmic*, *non-uniform rhythmic* and *random* patterns described above had been used in prior studies by our team (Chanes et al., 2013, 2015; Quentin et al., 2015; Stengel et al., 2021; Vernet et al., 2019). For all TMS patterns, the last pulse was delivered 33 ms before the onset of the visual target on the computer screen.

Each TMS pattern was tested in separate experimental blocks in which active and sham trials were randomly interleaved (50% active and 50% sham TMS trials). Participants performed 2 experimental sessions on two separate days (with an interval of a least 72 hours and a maximum of 7 days between sessions) to avoid carry-over effects and accrue sufficient EEG datasets for each TMS condition. Experimental procedures were identical in both sessions. Following a titration procedure for target contrast and after a block of task familiarization and training, participants carried out 4 experimental blocks in which each of the four TMS patterns presented above was tested. The order of TMS blocks for the two experimental sessions was counterbalanced across participants to avoid order-biases effects.

TMS stimulation intensity was fixed for all participants at a level of 55% maximal simulator output. Note that intensity was not adapted to each individual’s resting motor threshold (RMT). This is because motor cortex excitability was shown to predict very poorly excitability estimates in other cortical areas (Kähkönen et al. 2005; Stewart et al. 2001). Additionally, this same TMS intensity level (corrected to compensate EEG electrode thickness) has induced, at the group level, signs of local entrainment, increases of inter-regional connectivity and impacted visual perception outcomes (Stengel et al., 2021; Vernet et al., 2019). In any case, to allow across-study comparisons, at the end of each session the individual RMT of hand motor responses were determined on the *abductor pollicis brevis* (APB) muscle of each participant for the left primary motor (M1) cortex, and documented as the TMS intensity yielding right thumb activation in about 50% of stimulation attempts (Rossini et al., 2015). The average RMT of our cohort of participants was 66 ± 9% (mean ± SD) of the maximum stimulator output and our fixed TMS intensity translated into a stimulation intensity of 83 ± 12% (mean ± SD) of their individual motor thresholds.

### EEG recordings

EEG signals were recorded concurrently with stimulation pulses by means of 60 scalp electrodes connected to two 32-channel long-range TMS compatible amplifiers (BrainAmp DC and BrainVision Recording Software, BrainProducts GmbH). TMS-compatible passive EEG electrodes (3 mm thick) mounted on a cap centered at the head vertex (Cz) were located on specific scalp sites according to the international 10-20 EEG system. A taped electrode on the tip of the nose served as a reference, whereas a ground lead was placed on the right earlobe. EOG signals were recorded from 4 additional electrodes positioned on the left and right temples and above and below the left eye. Electrode impedances were monitored and kept below 5 kOhm throughout all sessions. EEG signals were digitized at a sampling rate of 5 kHz.

### EEG artifact removal procedure

EEG signals were analyzed with the FieldTrip toolbox (Oostenveld et al., 2011) running on MATLAB R2017b. EEG and EOG data were first epoched in a [-2 2] seconds window centered (t=0) on the onset time of the visual target. EEG traces from trials in which central fixation requirements were violated were automatically excluded. Additionally, the onset time of TTL trigger signals commanding the delivery of TMS pulses were automatically verified for each trial and rare events for which onset time proved temporally imprecise were excluded. Following visual inspection of all trials, those containing blinks and muscle artifacts were also eliminated from further analyses. After exclusions, an average number of 121 ± 14 (mean ± SD) viable trials per each TMS condition was retained for further steps

Expectedly, the electromagnetic field generated by brief TMS pulses induced a high amplitude electrical artifact on our EEG signals which had to be removed from our datasets. To this end, data across a [-4, +12] ms window centered on the onset of each TMS pulse was removed and ‘blank’ EEG epochs were filled with a shape-preserving piecewise cubic interpolation. To avoid any biases, the exact same artifact removal and interpolation procedure was applied to the EEG epochs of sham TMS trials. Once artifacts were removed, EEG signals were down-sampled to 500 Hz. Trials from all experimental conditions (i.e., active/sham trials for each of the 4 TMS tested patterns types) within each recording session were collected into two separate datasets. Then, two separate Independent Component Analyses (ICAs) corresponding to each experimental session were performed on the data. Such analysis did not artificially introduce any biases since trials gathered across all experimental conditions (4 TMS sham or active patterns) underwent the same ICA. Artefact components were identified based on the guidelines by Rogasch et al. (2014). This procedure enabled the removal of residual TMS artifacts lasting longer than 12 ms, which remained in EEG epochs after artifact cleaning and data interpolation processing steps. Components associated to eye movements, electrode malfunction and 50 Hz power line artifacts were also eliminated. An average of 9 ± 3 (mean± SD) components (out of 60 total components) were removed from each individual dataset.

Once the signal was calculated back to the electrode level, cleaned EEG datasets were separated into the following eight TMS experimental conditions: active/sham *rhythmic* TMS, active/sham *non-uniform rhythmic* TMS, active/sham *random* TMS and active/sham *irregular* TMS. Datasets evaluating the same TMS experimental condition from the two experimental sessions were combined for a common analysis.

### EEG outcome measures and analyses

EEG signals were transformed into the time-frequency domain with a 3-cycle Morlet wavelet analysis computed on a [-500 +500] ms window (centered on visual target onset, t=0) for frequencies between 6 and 50 Hz. In the time-frequency domain, we calculated measures of Power and Inter-Trial Coherence (ITC). ITC quantifies the level of oscillatory phase alignment across trials by averaging signal phase at each time-frequency point over all trials. Power was expressed in decibels (dB) relative to a baseline period of 2 oscillation cycles prior to the central alerting cue onset (i.e., the central cross becoming larger and preceding visual target onset by 233 ms).

First, we focused our analysis on a frequency band ([25 35] Hz) and time window of interest ([-133 0], t=0 visual target onset) and calculated the topographies for power and ITC across all the electrodes of our EEG grid. The frequency band of interest was centered on the 30 Hz frequency of our *rhythmic* TMS pattern and the time period of analyses included the 100 ms length of our 4 pulse TMS bursts. Second, we concentrated our analysis on a group of electrodes of interest (F1, F3, FC1, FC3 according to the 10-20 international EEG system) and calculated power and ITC over the complete time-frequency space. These specific scalp electrodes were selected since they were the closest to the center of the TMS coil targeting the left FEF.

Next, we aimed at evaluating the level of ‘noise’ of an EEG signal, a measure which depends on the level of predictability of the signal. Indeed, a completely regular and predictable signal, such as for example, a pure sinusoidal oscillation, has null ‘noise’ levels. A signal with several embedded frequencies, hence more heterogeneous and with broader band power spectrum, is more irregular or, in other words, more unpredictable and ultimately ‘noisier’ than the former. For instance, white noise, which is a completely random signal, has a flat power spectrum (i.e., equal contribution of all frequency bands) is totally unpredictable and considered very ‘noisy’.

Three main measures were used in the current study to estimate the level of internal neural ‘noise’ present in brain activity according to EEG signals recorded under the influence of our four different TMS stimulation patterns: Power peak-width, Sample Entropy (SE) and Multi-Scale Entropy (MSE) (Fig. 2).

**Figure 2:**
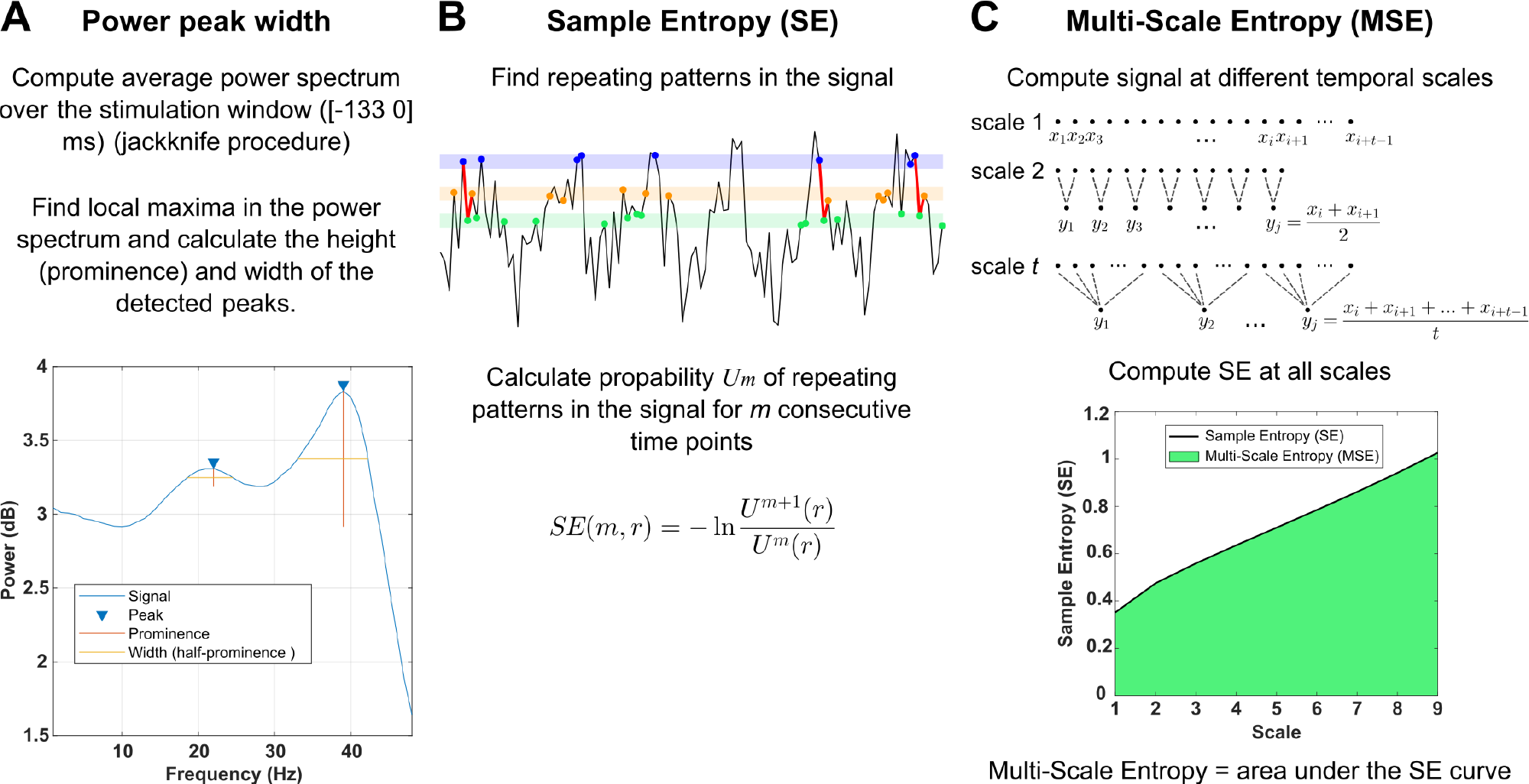
Outcome measures used to evaluate neural noise levels induced by TMS patterns in EEG signals. **(A)** Width and number of peaks in the power spectrum of EEG time series within a time window of interest including TMS delivery. Peaks are local maxima in the power spectrum. Peak width is calculated at the half-prominence. Peak prominence is the difference between the local maximum and the smallest local minimum between this local maximum and the next higher local maximum. **(B)** Sample Entropy is estimated in a time series by counting repeating patterns of length *m* and *m+1* in the signal. In the middle panel, all dots labelled in blue are within a distance *r* of each other, the same applies to dots labelled in green and orange. A repeating motif (in the example represented, a sequence of blue, green and orange dots) is identified when the same sequence (blue, green, orange) re-appears later in the signal (repeated motifs are marked in red). A ratio of probability of repeating patterns of length *m* and repeating patterns of length *m+1* is then computed as an estimation of the entropy of the signal. **(C)** Multi-Scale Entropy estimates changes of Sample Entropy across several time scales. Time scales are estimated by averaging the signal inside non-overlapping time windows of varying length (upper panel). The shape of the Sample Entropy curve across time scales (lower panel, black line) informs on the Multi-Scale Entropy content of the signal. A single, lower dimensionality value of Multi-Scale Entropy was estimated by calculating the area under the Sample Entropy curve (lower panel, green area).

In the first of our approaches, we calculated the peak-width of the frequency band of oscillations enhanced during TMS (Fig. 2A). Such measure, which informs on the heterogeneity of the signals or, in other words, the variety of oscillation frequencies carried by EEG signals, was employed as a proxy of ‘noise’ (hence ‘unpredictability’). Accordingly, the ‘nosier’ a signal, the less frequency-tuned, the broader peak-width and vice-versa. To this end, we averaged the power spectrum over frequencies [6 45] Hz across the stimulation time window ([-133 0] ms, between the 1st pulse of each burst and visual target onset, t=0). We then detected local maxima in the average power spectrum. The width of each local maximum, or peak, in the signal was calculated at half prominence. The prominence or height of a peak was determined relative to the lowest local minimum between a given peak and the next peak higher than the current one (Fig. 2A, bottom panel).

Given the low signal-to-noise ratio of scalp EEG signals, reliable peaks could not be identified on individual datasets (Kiesel et al., 2008; Ulrich & Miller, 2001). Therefore, we calculated the average power spectrum during TMS stimulation on grand averages across participants. In order to estimate the variance in the measure of peak-width for our group of participants we applied the jackknife procedure, which, for a sample of N participants, computes for each *i* (*i*=1, …, N) the grand average signal over a subsample of N-1 participants by omitting participant *i* in the dataset. Peaks in the power spectrum and their width were estimated for each of the N subsampled grand averages. Since such measures are based on grand averages over a pool of participants and thus vary at each iteration only by a single individual, estimations made with this method have a very low error variance, therefore the standard error has to be corrected according to the following formula:

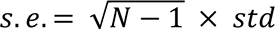

Where *s.e*. is the corrected standard error and *std* corresponds to the standard deviation of the jackknife subsampled measures. To test for significance, t- and F-statistics also had to be corrected for the reduced error variance in the following way (Ulrich & Miller, 2001):

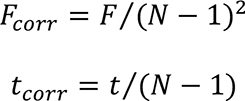

An identical analysis was conducted on ITC peak-width (see Supplementary Results).

In our second approach, we computed, in the time-domain, the Sample Entropy (SE), a direct measure of entropy which is a metric of disorder of the contents of a system, and Multi-Scale Entropy (MSE), which is based on the former but directly evaluates signal complexity (Costa et al., 2002, 2005). Indeed, a measure of ‘complexity’ differs from a measure of ‘entropy’ or ‘noise’ in that neither a completely regular or predictable signal, such as a pure oscillation (with null or very low entropy), nor, paradoxically, a completely random and unpredictable signal (with maximal entropy levels) such as white noise, exhibit high complexity. Indeed, both signals contain very poor information because they are governed by very simple laws.

Sample Entropy (SE) identifies repeating patterns within a signal. Two time points are considered similar if they are within a distance *r* of each other and thus, for any pattern of *m* consecutive time points [u_1_, u_2_, …, u_m_] in the signal another sequence [v_1_, v_2_, …, v_m_] is considered a repetition of the first pattern if |v_1_ – u_1_| ≤ *r*, |v_2_ – u_2_| ≤ *r*, …,|v_m_ – u_m_| ≤ *r* respectively. In other words, a sequence of consecutive time points is considered a repetition of a pattern if each time-point in the sequence is within a distance *r* from the corresponding time point in the pattern (Fig. 2B, sequences marked in red are repeated). The search for repeating patterns in the signal is done for each sequence of *m* consecutive time points in the data (excluding self-matches) and yields the probability U^m^(r) that two sequences of *m* time points are within a distance *r* of each other. The same calculation can be performed for patterns of *m+1* time points. Sample Entropy (SE) is then defined as:

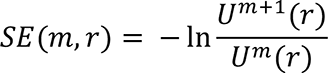

This measure represents the conditional probability that, knowing that a pattern is repeated for *m* consecutive time points, it will also be repeated for *m+1* time points. Essentially, SE evaluates the probability that the *m+1* time point can be predicted when following a known pattern of *m* time points. The lower this probability, the less predictable a signal is and the higher its entropy.

Signal complexity is evaluated by repeating the measure of Sample Entropy at several time scales of a signal. The signal at time scale *t* is the original signal averaged inside non-overlapping time windows of length *t* (Fig. 2C, upper panel). In this context, signal complexity, as evaluated by Multi-Scale Entropy, is represented as the evolution of the Sample Entropy across time scales.

Accordingly, a very predictable signal will have low Sample Entropy values at all scales. Additionally, a signal such as white noise, that is very unpredictable, will have very high Sample Entropy at low time scales however, when long stretches of data are averaged for higher time scales, the random signal that is white noise will average to a constant signal at 0 and will exhibit very low values of Sample Entropy. Only complex time series that exhibit, information-wise, a very rich and unpredictable signal over varied temporal scales will show high Sample Entropy values at all time scales. Therefore, a signal that exhibits higher Sample Entropy values at a majority of time scales compared to another signal can be considered relatively more complex (Costa et al., 2005).

In our study, to reduce the dimensionality of the Multi-Scale Entropy value and compare signal complexity estimates over the whole EEG electrode array across different TMS patterns, we calculated the area under the profile of Sample Entropy values along time-scales. We accepted the assumption that a higher area under the Sample Entropy profile of EEG signals reflected higher Sample Entropy at a majority of time scales and therefore indicated a higher degree of signal complexity. We referred to the area under the Sample Entropy profile as the Multi-Scale Entropy (MSE) measure (Fig. 2C, lower panel).

We calculated Sample Entropy and Multi-Scale Entropy from EEG epochs during the stimulation time window [-133 0] ms (between the 1st TMS pulse and visual target onset, t=0) for each EEG grid electrode. Considering that values of Sample Entropy at each time scale are more reliable for longer signals we computed Multi-Scale Entropy on EEG epochs recorded at 5 kHz sampling rate, hence with the highest number of available data points. We set Sample Entropy parameters at m=2 and *r* as 15% of the signal standard deviation (Costa et al., 2005). Since EEG epochs associated to TMS stimulation were too short to yield reliable values of Sample Entropy at the higher scales (non-finite values of Sample Entropy), analyses presented in the current study included up to 9 scales (even if up to 14 scales were calculated).

### EEG statistical analyses

For both topographical and time-frequency maps of power, ITC and MSE, we performed comparisons between active and sham trials for each TMS patterns as well as direct two-by-two comparisons between active trials across the 4 TMS pattern types: *rhythmic*, *non-uniform rhythmic*, *random* and *irregular*. Each pair was compared with two-tailed paired Student’s *t*-tests. Comparisons between topographical maps were performed for each EEG electrode whereas those between time-frequency maps were calculated for each time-frequency point within [6 45] Hz and across a time window of [-300 200] ms (visual target onset t=0). Cluster-based permutation tests, used to correct for multiple comparisons, were performed on clusters of neighboring electrodes or time-frequency points that exceeded the significance threshold (alpha = 0.01) in paired t-tests and assigned a statistic which was the sum of T-statistics of each point of the cluster.

A non-parametric permutation test (10000 permutations, Montecarlo sampling method) was applied to cluster statistics. The statistical analyses displayed in figures 3, 4, 5, 6 and 8 show clusters that exceeded the threshold for significance (alpha=0.05) for the two-sided permutation test. Full details on the statistical results (including cluster extents, T-statistic and p-value) are provided in the Supplementary Materials (see Tables S1, S2, S3, S4 and S5). Any effect observed on any EEG signal is likely to last over several time points and spread over adjacent electrodes, hence cluster-based permutations is a highly sensitive method to correct for multiple comparisons in this type of data because it is adapted to a high degree of correlation in time and space (Maris & Oostenveld, 2007). However, there is currently no consensus on how to apply cluster-based permutations to interaction effects for ANOVA analyses (Edgington & Onghena, 2007; Suckling & Bullmore, 2004), therefore pairwise comparisons were performed between TMS experimental conditions.

**Figure 3.**
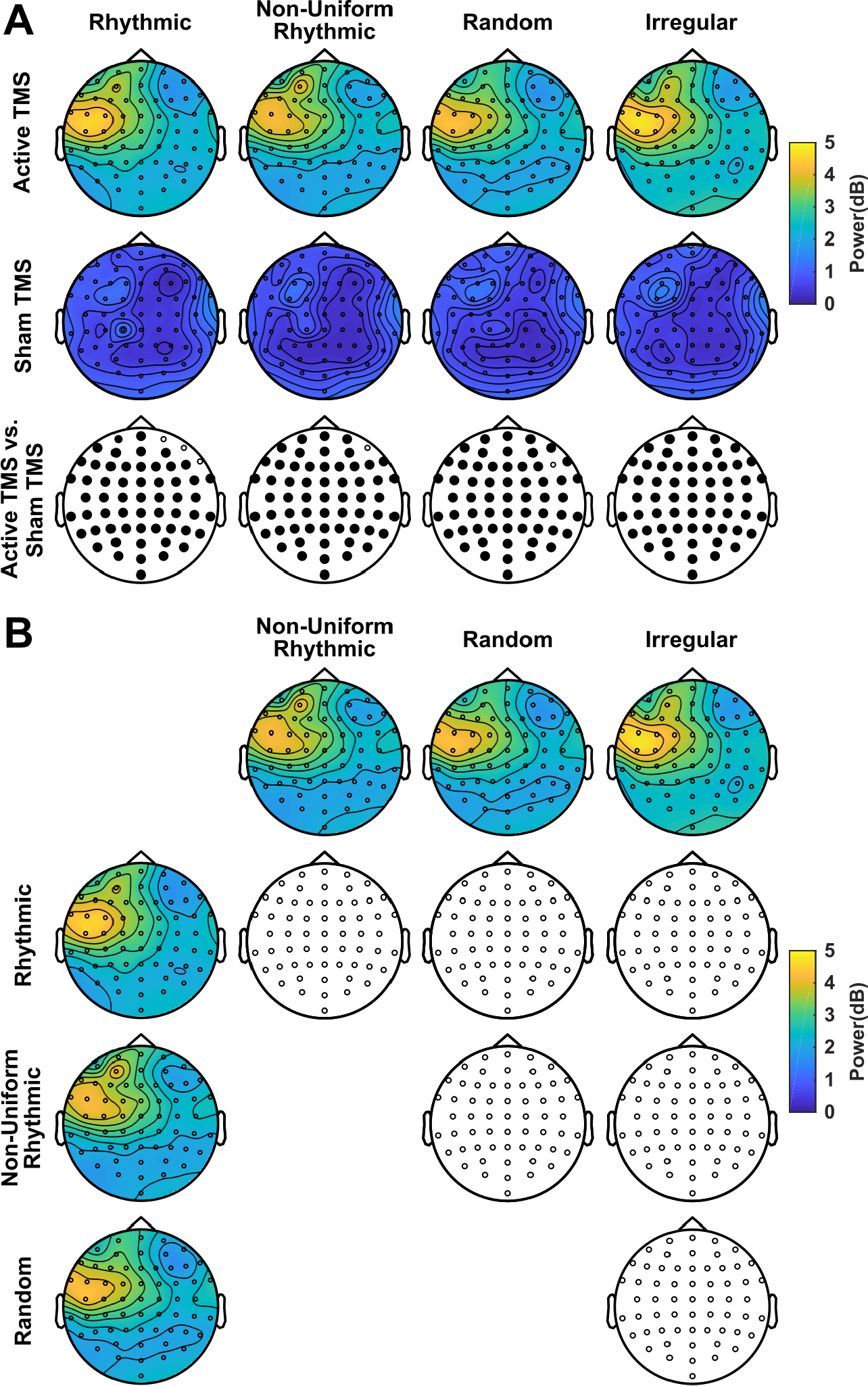
Impact of rhythmic and non frequency-specific TMS patterns on left frontal high-beta oscillation power. Topographical maps representing data in the [25 35] Hz frequency band during stimulation (time window [-133 0] ms centered on visual target onset). **(A)** Comparisons of high-beta power between active (first row) and sham (second row) TMS bursts for each pattern type (*30 Hz rhythmic*, and 3 *non frequency-specific* patterns: *non-uniform rhythmic*, *random* and *irregular*). The bottom row shows the outcomes of pairwise (active vs. sham TMS) cluster-based statistical permutation tests. Bolded electrodes represent clusters of contacts that reached statistical significance (p<0.05). **(B)** Direct two-by-two comparisons of high-beta Power between the different active TMS patterns. Colored maps represent distribution of Power over the scalp for all four TMS patterns (*30 Hz rhythmic*, *non-uniform rhythmic*, *random* and *irregular*). Uncolored maps display the results of the cluster-based statistical permutation tests for the pairwise comparison of active trials in the two topographical maps represented at the top of the column and the left of the row. Notice that no EEG contacts showed statistically significant differences (p>0.05) in any of the comparisons. Both *30 Hz rhythmic* TMS and the three *non frequency-specific* TMS patterns increased amplitude of high-beta oscillations over the whole scalp during active stimulation compared to sham.

Statistical analyses on the peak-width for power increases between our 4 active TMS patterns were carried out with a one-way repeated measures ANOVA with factor *TMS pattern* (*rhythmic*, *random* and *irregular*). The *non-uniform rhythmic* pattern had to be excluded from such analysis because its power spectrum exhibited two frequency peaks and therefore could not be fairly compared in terms of level of EEG ‘noise’ with the remaining 3 patterns which featured a single frequency peak in their power spectrum.

## Results

### Impact of non frequency-specific TMS patterns on high-beta oscillations

We first tested the impact of *30 Hz rhythmic* and the three *non frequency-specific* TMS patterns *(non-uniform rhythmic*, *random* and *irregular*) on highly predictable and regular high-beta oscillations. To this end, we examined modulations of high-beta power ([25 35] Hz) for all scalp electrodes during the stimulation window ([-133 0] ms centered on target onset, t=0) (Fig. 3).

Our analyses revealed that, as expected, active stimulation with 30 Hz patterns significantly increased high-beta power compared to sham stimulation. Surprisingly however, the three active *non frequency-specific* bursts also showed significant increases of high-beta power, even if they did not contain 30 Hz motifs embedded in their temporal structure (Fig. 3A) Moreover, direct two-by-two comparisons of active TMS conditions failed to reveal significant differences in high-beta power during stimulation between *rhythmic*, *non-uniform rhythmic*, *random* and *irregular* active TMS patterns, suggesting that all four TMS patterns did not differ with regards to their ability to entrain 30 Hz oscillations (Fig. 3B). See supplementary table S1 for full details on the statistical results for all comparisons.

We also analyzed the degree of phase-locking for high-beta activity during stimulation by calculating inter-trial coherence (ITC) at this specific frequency band (25-35Hz). As reported above for oscillation power, a comparison between active and sham TMS patterns revealed significant increases of high-beta phase-locking for all EEG grid electrodes during the 30 Hz *rhythmic* and also for the three *non frequency-specific* patterns tested in our study (Fig. 4A). However, this time, direct comparisons between TMS patterns showed that not all patterns demonstrated statistically equivalent effects on ITC. Instead, the *random* pattern showed significantly lower high-beta ITC than active 30 Hz *rhythmic*, *non-uniform rhythmic* and *irregular* bursts. Additionally, two isolated grid electrodes (FCz and F2) displayed significantly higher ITC levels of for active *irregular* patterns compared to the active 30 Hz *rhythmic* condition (Fig. 4B). See supplementary table S2 for full details on the statistical outcomes for all comparisons.

**Figure 4.**
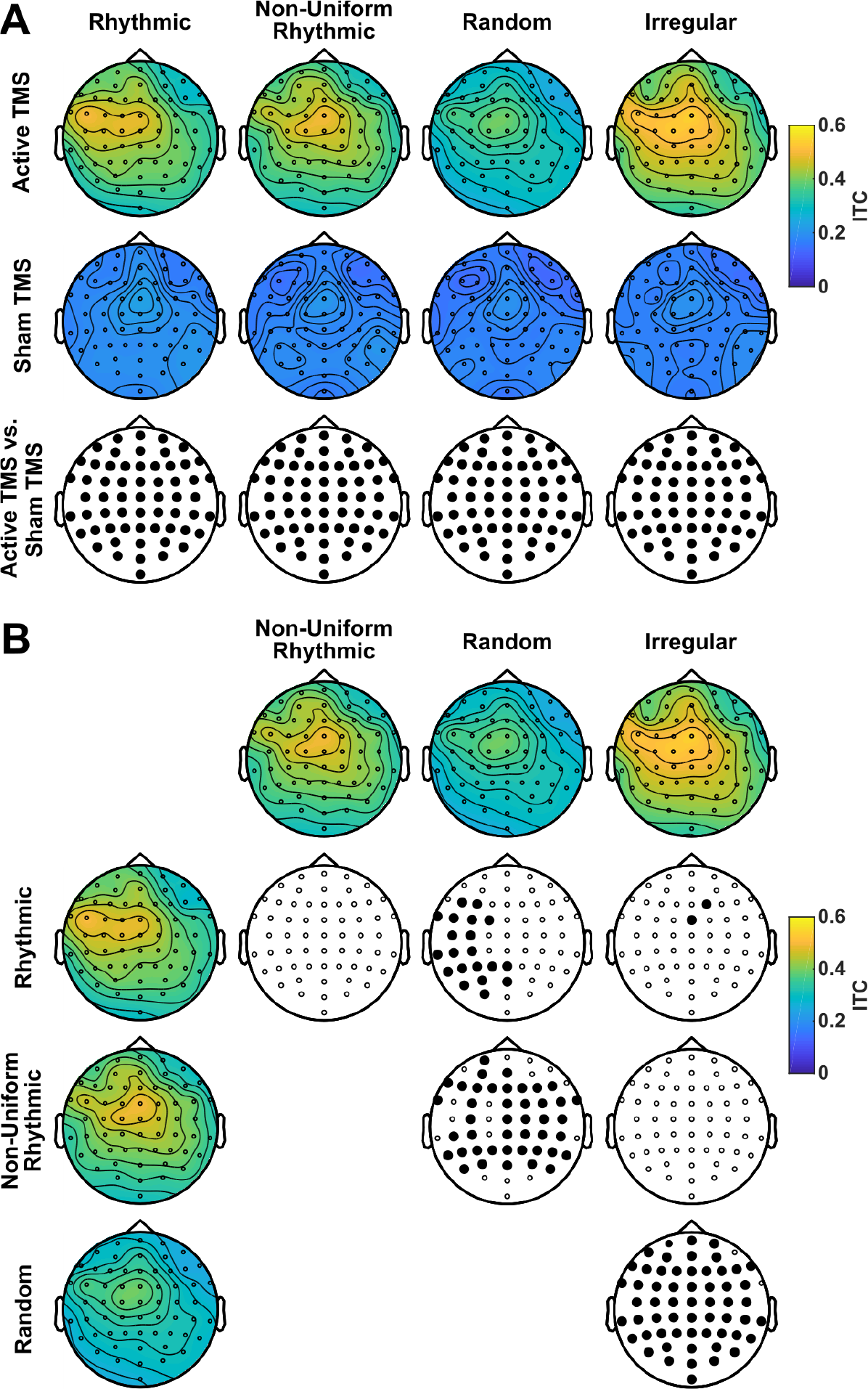
Impact of rhythmic and non frequency-specific TMS patterns on left frontal high-beta oscillatory phase alignment. Topographical maps representing data in the [25 35] Hz frequency band during stimulation (time window [-133 0] ms centered on visual target onset). **(A)** Comparisons of inter-trial coherence (ITC) between active (first row) and sham (second row) TMS bursts for each pattern type (*30 Hz rhythmic*, and 3 *non frequency-specific* patterns: *non-uniform rhythmic*, *random* and *irregular*). The bottom row shows the outcomes of pairwise (active vs. sham TMS) cluster-based statistical permutation tests. Bolded electrodes represent clusters of contacts that reached statistical significance (p<0.05). **(B)** Direct two-by-two comparisons of ITC levels between the different active TMS patterns. Colored maps represent distribution of ITC over the scalp for all four TMS patterns (*30 Hz rhythmic*, *non-uniform rhythmic*, *random* and *irregular*). Uncolored maps display the results of the cluster-based statistical permutation tests for the pairwise comparison of active trials in the two topographical maps represented at the top of the column and the left of the row. Bolded EEG electrodes represent EEG contacts or clusters of EEG contacts for which differences reached statistical significance (p<0.05). Both *30 Hz rhythmic* TMS and the three *non frequency-specific* TMS patterns increased phase-alignment of high-beta oscillations over the whole scalp during active stimulation compared to sham. Direct pairwise comparisons of ITC between active TMS patterns revealed that *random* TMS achieved lower increases of high-beta inter-trial phase-locking than the other three active TMS patterns.

Taken together, these analyses suggest that the *non frequency-specific* TMS patterns, designed not to contain high-beta rhythms in their temporal structure were all able to increase high-beta power and, for a majority of stimulation conditions, also high-beta ITC as strongly as a pure 30 Hz *rhythmic* TMS stimulation (specifically designed to entrain high-beta cortical oscillations) did.

### Frequency-specificity of oscillatory entrainment by TMS patterns

We then explored the hypothesis that the three *non frequency-specific* TMS patterns *(non-uniform rhythmic*, *random* and *irregular*) increased oscillations across a broader frequency band than 30 Hz *rhythmic* TMS patterns. This outcome would contribute to indirectly substantiate their ability to entrain less predictable and more complex, hence more ‘noisy’, internal neural activity in left frontal systems. To this end, for a group of selected leads (scalp electrodes F1, F3, FC1 and FC3) located in the vicinity of the targeted left FEF region, we examined modulations of power and ITC over a broader frequency range ([6 45] Hz). Additionally, we assessed power and ITC over a longer time window ([-500 500] ms centered on visual target onset) to test if oscillatory modulations were restricted to the time period of stimulation.

Comparisons between active TMS patterns and their associated sham conditions revealed, during the delivery of 30 Hz *rhythmic* patterns and also for the three *non frequency-specific* bursts, increased oscillation power (Fig. 5A) and ITC (Fig. 6A) in a broad frequency band, which was not strictly limited to 30 Hz. Direct two-by-two comparison of oscillation power during active stimulation showed no significant differences in the frequency bands modulated by the 3 *non frequency-specific* patterns compared to the *rhythmic* pattern (Fig. 5B, first row, signal between the two vertical red dotted lines indicating the stimulation window). However, direct two-by-two comparisons of ITC showed that compared to the active *rhythmic* TMS condition, active *irregular* TMS increased phase-locking across a band extending to the low-beta range (12-20 Hz) (Fig. 6B).

**Figure 5.**
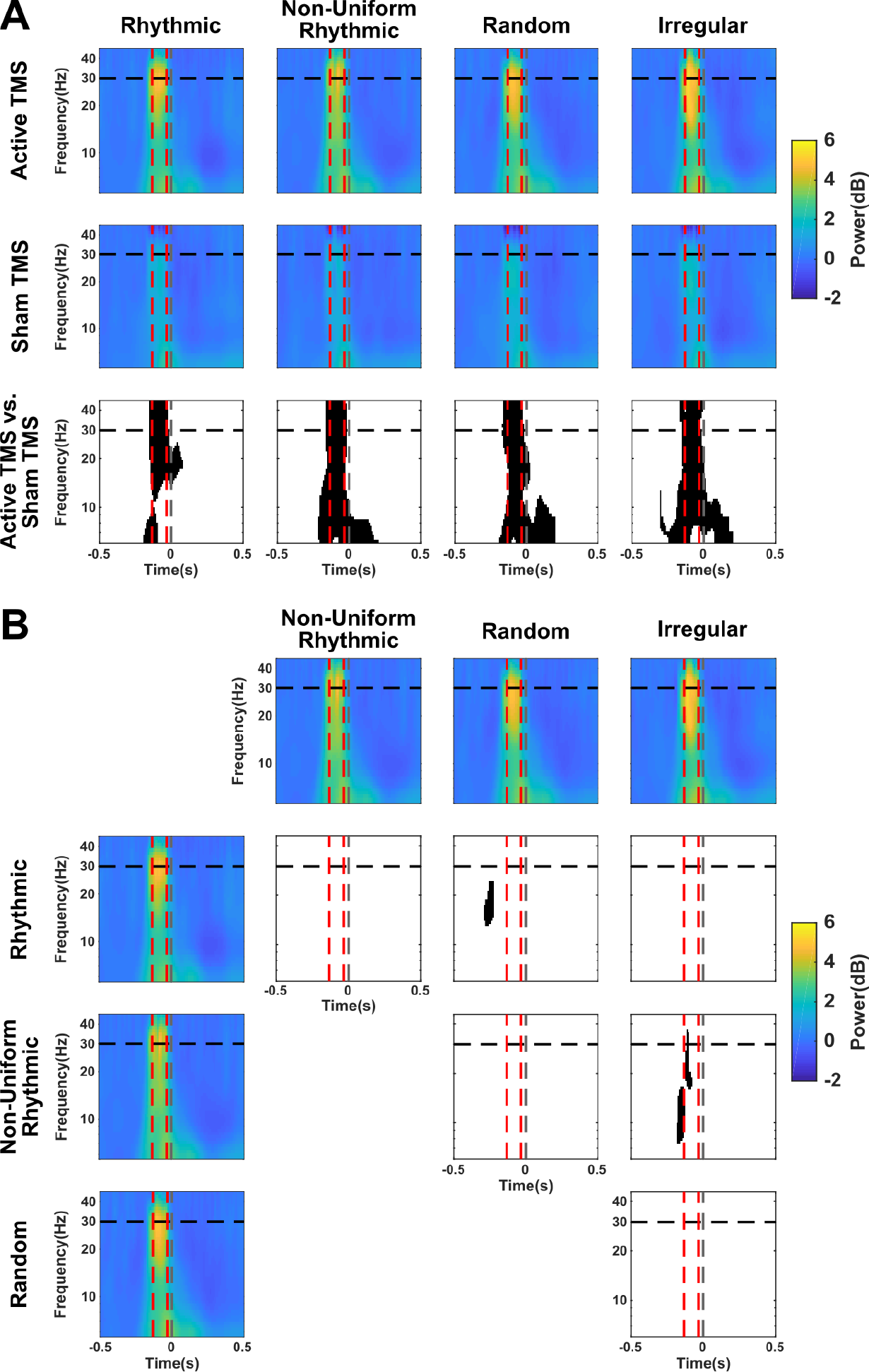
Frequency-specific modulation of cortical oscillations power by rhythmic and non-frequency-specific TMS patterns in left frontal regions. Time-frequency maps in a cluster of left frontal electrodes (F1, F3, FC1, FC3) closest to the center of the stimulation coil. Time is centered on the onset of the visual target (dotted grey vertical line). The two red dotted vertical lines signal the first (−133 ms) and last (−33 ms) TMS pulses of the burst. The black dotted horizontal line indicates the frequency (30 Hz) of the *rhythmic* TMS pattern. **(A)** Comparisons of high-beta power between active (first row) and sham (second row) TMS for each stimulation pattern (30 Hz *rhythmic*, and three *non frequency-specific* patterns: *non-uniform rhythmic*, *random* and *irregular*). The bottom row shows the results of pairwise (active vs. sham TMS) cluster-based statistical permutation tests. Black clusters indicate time-frequency points for which comparisons reached statistical significance (p<0.05). **(B)** Direct two-by-two comparisons across the 4 types of active TMS patterns. Colored maps represent time-frequency maps of power for all TMS patterns (30 Hz *rhythmic*, *non-uniform rhythmic*, *random* and *irregular*). Black and white maps show the outcomes of the cluster-based statistical permutation tests for pairwise comparisons between active TMS trials for the two time-frequency maps represented at the top of the column and the left of the row. Black clusters indicate time-frequency points for which comparisons reached statistical significance (p<0.05). Both *30 rhythmic* TMS and the 3 *non frequency-specific* TMS patterns increased amplitude of cortical oscillations over a wide frequency band during active compared to sham stimulation.

**Figure 6.**
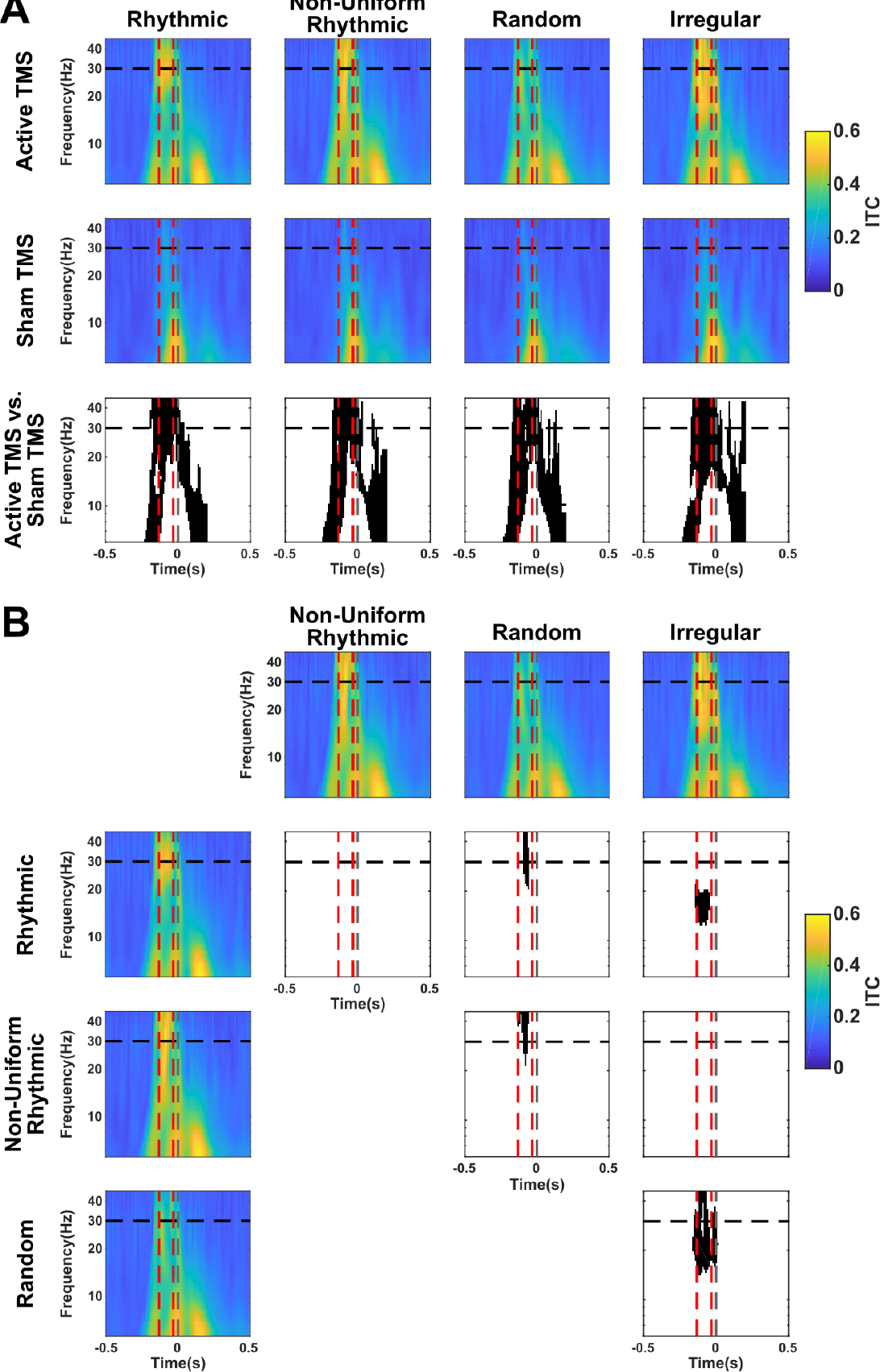
Frequency-specific modulation of oscillatory phase-alignment by rhythmic and non-frequency-specific TMS patterns in left frontal regions. Time-frequency maps in a cluster of left frontal electrodes (F1, F3, FC1, FC3) closest to the center of the stimulation coil. Time is centered on the onset of the visual target (dotted grey vertical line). The two red dotted vertical lines signal the first (−133 ms) and last (−33 ms) TMS pulses of the burst. The black dotted horizontal line indicates the frequency (30 Hz) of the rhythmic TMS pattern. **(A)** Comparisons of inter-trial coherence (ITC) between active (first row) and sham (second row) TMS for each stimulation pattern (30 Hz *rhythmic*, and three *non frequency-specific* patterns: *non-uniform rhythmic*, *random* and *irregular*). The bottom row shows the results of pairwise (active vs. sham TMS) cluster-based statistical permutation tests. Black clusters indicate time-frequency points for which comparisons reached statistical significance (p<0.05). **(B)** Direct two-by-two comparisons across the 4 types of active TMS patterns. Colored maps represent time-frequency maps of ITC for all TMS patterns (30 Hz *rhythmic*, *non-uniform rhythmic*, *random* and *irregular*). Black and white maps show the outcomes of the cluster-based statistical permutation tests for pairwise comparisons between active TMS trials for the two time-frequency maps represented at the top of the column and the left of the row. Black clusters indicate time-frequency points for which comparisons reached statistical significance (p<0.05). Both *30 rhythmic* TMS and the 3 *non frequency-specific* TMS patterns increased phase-alignment of cortical oscillations over a wide frequency band during active compared to sham stimulation. However, the active *irregular* TMS patterns achieved higher increases of oscillatory phase-locking in the low-beta band than the active 30 Hz *rhythmic* TMS pattern. Also note that the *random* TMS pattern phase-locked cortical oscillations trial-to-trial to a significantly lower degree than the remaining active TMS patterns.

We also noticed that, similarly to what was shown on the topography (Fig. 4B), active *random* bursts increased high-beta (∼30 Hz) phase-locking to a lesser degree than *rhythmic*, *non-uniform rhythmic* or *irregular* TMS patterns (Fig. 6B). For full details on the statistical outcomes reported in this section refer to supplementary tables S3 and S4.

In order to specifically estimate the heterogeneity of the entrained oscillations, which we used as a proxy for changes in the level of internal neural noise, we quantified the width of the frequency band showing power increases caused by TMS stimulation. A jackknife procedure was used to identify peaks on the grand-average power spectrum over the whole stimulation window ([−133 0] ms, t=0 ms corresponding to visual target onset) and the width of these peaks was extracted (Fig. 7A). Notice that for active *rhythmic*, *random* and *irregular* patterns, a single high-beta frequency peak (at ∼28 Hz, ∼27 Hz and ∼28 Hz, respectively) was identified in the power spectrum. Nonetheless, for *non-uniform rhythmic* TMS, in addition to the peak reliably identified at a high-beta frequency (∼31 Hz), 12 out of 15 iterations of the jackknife procedure also revealed a second peak, this time in the low-beta range (∼15 Hz).

**Figure 7.**
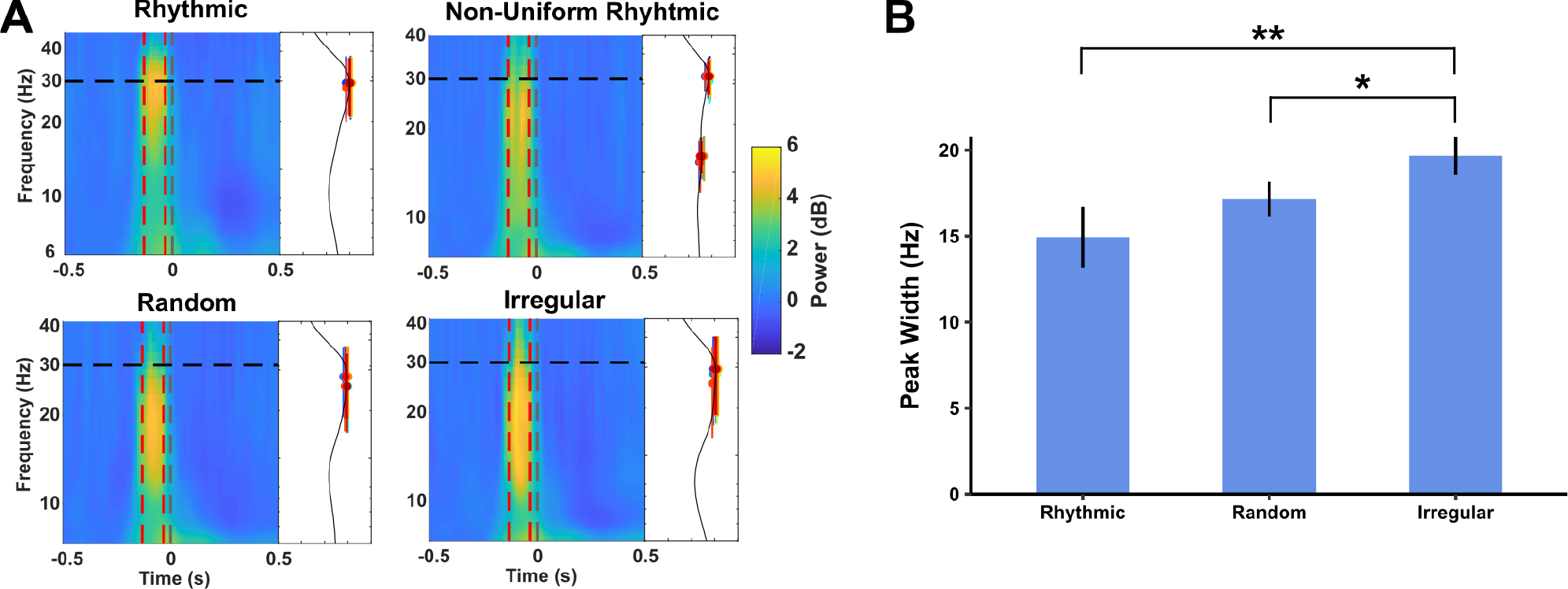
Width of power peak increases during 30 Hz rhythmic TMS compared to the three non frequency-specific TMS patterns delivered in a left frontal region. **(A)** Time-frequency maps in a cluster of left frontal electrodes (F1, F3, FC1, FC3) closest to the center of the stimulation coil during active stimulation trials represented for each stimulation pattern (30 Hz *rhythmic*, and three *non frequency-specific* patterns: *non-uniform rhythmic*, *random* and *irregular*). The two red dotted vertical lines signal the first (−133 ms) and last (−33 ms) TMS pulses of the burst. The black dotted horizontal line indicates the frequency (30 Hz) of the rhythmic TMS pattern. Right marginal graphs of each time-frequency panel display the average power spectrum over the whole window of TMS delivery. Colored lines show the width of the peaks of oscillations power as detected with a jackknife procedure. **(B)** Comparison of width of peaks of power increase (expressed in Hz) during active stimulation between uniform *rhythmic* and *non frequency specific* (*random* and *irregular*) TMS. The so called *non-uniform rhythmic* TMS pattern is not presented in the figure, because its power spectrum during active TMS revealed two distinct peaks (∼30 Hz and ∼15 Hz, panel A) and such outcomes could not be directly compared to those displaying only a single peak in the high-beta range (∼30 Hz). Notice that error bars represent the standard error corrected for the reduced error variance obtained from the jackknife procedure. One-way ANOVA analysis yielded a significant main effect of TMS pattern. Notice that *irregular* TMS patterns increased cortical oscillations in a significantly wider frequency band than uniform *rhythmic* or *random* TMS patterns. Results of the post-hoc t-tests are indicated as follows: ** p < 0.01, * p < 0.05.

Peak-width values for power increases were compared with a one-way ANOVA. However, for the *non-uniform rhythmic* TMS condition, the presence of two separate peaks prevented us from estimating a single value of power peak-width. Therefore, we judged that this TMS condition could not be compared directly to the other three featuring power peaks at a single frequency band and, accordingly, it was not included in the ANOVA. Statistical results showed significant differences across active *rhythmic*, *random* and *irregular* stimulation patterns (F(2,41) =3.309, MSE=0.13, p<0.05, after correction for reduced error variance from the jackknife procedure). Planned two-tailed paired Student *t*-test showed that active *irregular* TMS increased oscillation power in a significantly broader band than 30 Hz *rhythmic* (T(14) = 2.68, p<0.01, corrected for reduced error variance by jackknife procedure) or *random* TMS bursts (T(14) = 2.016, p<0.05, corrected for reduced error variance by jackknife procedure) (Fig. 7B). An identical analysis performed on peak-width for ITC increases during stimulation yielded similar statistical results (see Supplementary Materials, Fig. S1).

This analysis suggests that *non frequency-specific* TMS patterns differed from uniform *rhythmic* stimulation with regards to the width of the frequency band they were able to modulate. *Non frequency-specific* TMS patterns increased cortical oscillations amplitude in a wider beta band or in two distinct low-beta and high-beta frequency bands.

### Modulation of signal complexity by non frequency-specific TMS activity

To confirm our conclusions on the modulation of internal neural noise levels contained in EEG signals by *non frequency-specific* TMS patterns, we turned to two more direct measures of ‘noise’ based this time on an assessment of the predictability and regularity of EEG signals during stimulation: Sample Entropy (SE) and Multi-Scale Entropy (MSE). Multi-Scale Entropy specifically estimates the complexity of a signal by calculating Sample Entropy at several time scales in which the signal can be decomposed. Both measures were computed in the time-domain on EEG epochs associated to the TMS delivery window ([−133 0] ms, t=0 visual target onset) for all grid electrodes.

The level of Sample Entropy raised gradually across time-scales (note steeper increases for active compared to sham TMS patterns) (Fig. 8A, here illustrated for electrodes F1, F3, FC1, FC3, located directly under the stimulation coil, but the same gradual increase is observed for all electrodes). This pattern of across-scale Sample Entropy characterizes a ‘complex’ signal which is irregular and non-predictable over multiple time-scales (Costa et al., 2002, 2005; Zhang, 1991). In order to reduce the dimensionality of our measure, for subsequent analyses we calculated for each participant, each stimulation condition and each electrode the area under the across-scale Sample Entropy profile and used this measure as an across-scale integrated estimate of Multi-Scale Entropy which we represented as EEG topographic maps (Fig. 8B).

**Figure 8.**
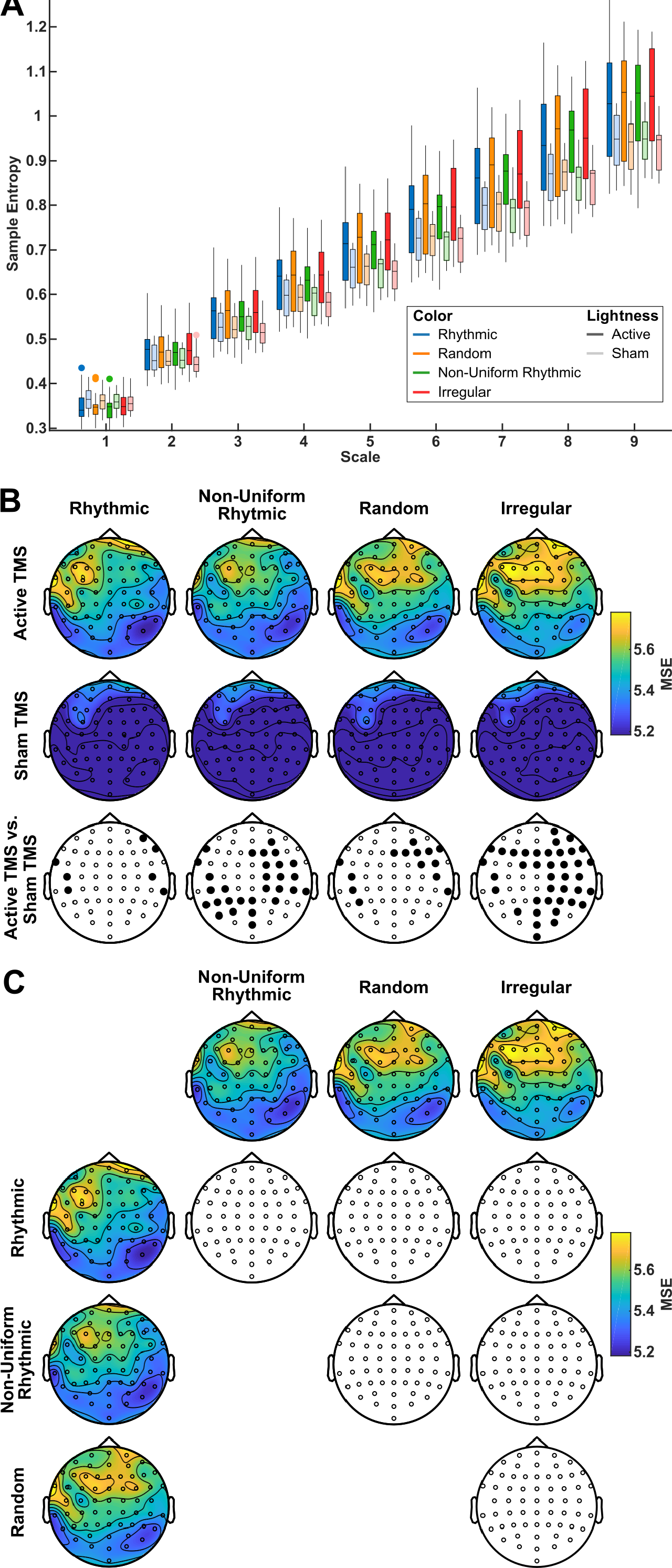
Modulation of EEG signal complexity by rhythmic and non frequency-specific TMS patterns in left frontal regions. **(A)** Bar plot of Sample Entropy (SE) across time scales for left frontal electrodes (F1, F3, FC1, FC3) closest to the center of the TMS stimulation coil. Box-plot color codes identify each of the 4 TMS pattern tested in the study (Blue: 30 Hz *rhythmic* TMS, and three non frequency-specific patterns, Orange: *random* TMS, Green: *non-uniform rhythmic* TMS, and Red: *irregular* TMS). The solid colors indicate bar plots for *active* TMS condition, whereas ‘pastel’ or less saturated colors identify box-plots associated to each of the sham TMS patterns. Notice that the estimated values of SE increases across time-scales for all TMS patterns in both active and sham TMS trials. This suggests that EEG time series contain a measure of unpredictability and noise at several time-scales, which is the hallmark of a complex signal. **(B)** Comparisons between active TMS (1st row) and sham TMS (2nd row) topographical maps of the areas under the curve of SE across time-scales, a measure entitled MSE (Multi-scale entropy), are shown for each TMS pattern (30 Hz *rhythmic*, and the 3 *non frequency-specific* patterns: *non-uniform rhythmic*, *random* and *irregular*). The bottom row shows the results of pairwise (active vs. sham TMS) cluster-based statistical permutation tests. Bolded EEG electrodes represent clusters of sensors that reached statistical significance (p<0.05). Non frequency-specific TMS increased MSE in clusters of left frontal and bilateral parietal EEG contacts. Comparison between active and sham *rhythmic* TMS pattern showed only sporadic significant differences in isolated EEG leads. **(C)** Direct two-by-two comparisons of MSE between the different active TMS patterns. Colored maps represent distribution of MSE over the scalp for all four TMS patterns (*30 Hz rhythmic*, *non-uniform rhythmic*, *random* and *irregular*). Uncolored maps display the results of the cluster-based statistical permutation tests for the pairwise comparison of active trials in the two topographical maps represented at the top of the column and the left of the row. Notice that no EEG contacts showed statistically significant differences (p>0.05) in any of the comparisons.

**Figure 9.**
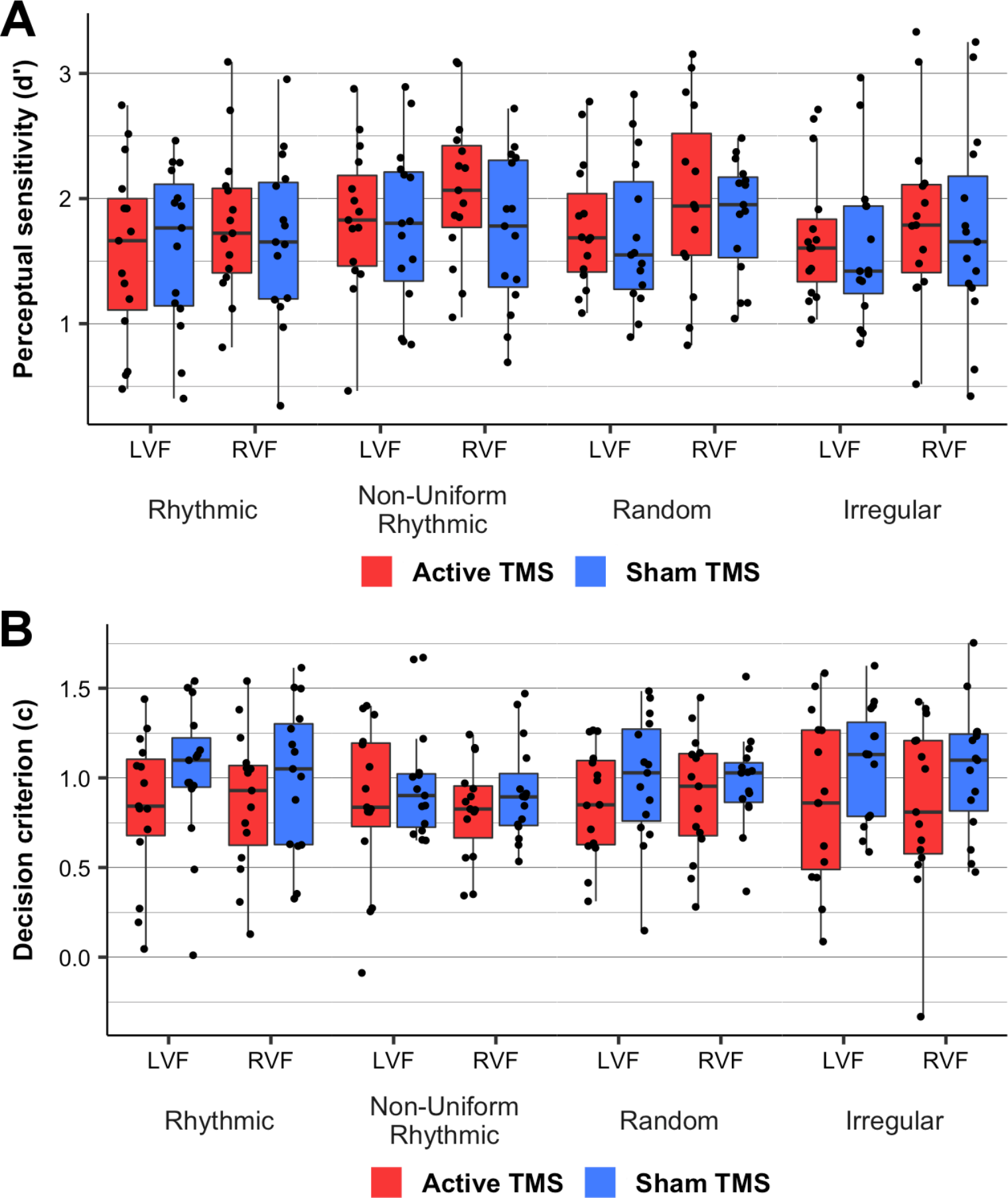
Impact of rhythmic and non frequency-specific left frontal TMS patterns on conscious visual detection performance. Modulation of **(A)** perceptual sensitivity (d’) and **(B)** decision criterion (c) by active TMS patterns (in red) compared to sham TMS patterns (in blue). Data is presented for each TMS pattern (30 Hz *rhythmic* TMS and the 3 *non frequency-specific* TMS patterns: *non-uniform rhythmic*, *random* and *irregular*) separately for targets displayed in the left and the right visual fields (LVF and RVF, respectively). Boxes are drawn from the 25^th^ to the 75^th^ percentile and the horizontal line marks the median. Whiskers are drawn between the minimum and maximum data points, excluding any outliers. Black dots indicate individual measures. Note that no specific effects of *rhythmic* or *non frequency-specific* TMS patterns were revealed by our analyses on d’ or decision criterion. Nonetheless, the delivery of any of these 4 active TMS patterns, regardless of their temporal structure, lowered decision criterion (significant main effect of TMS condition) rendering participants less conservative when having to decide if a near threshold target had been presented or not.

Comparisons between active vs. sham trials revealed that *non frequency-specific* TMS patterns increased Multiple Sample Entropy in clusters of EEG contacts located over fronto-parietal regions. More specifically, active *random* stimulation increased Multi-Scale Entropy in a small cluster of right frontal electrodes, whereas *non-uniform rhythmic* and *irregular* active stimulation showed a more widespread topography extending to parietal electrodes. Active *non-uniform rhythmic* stimulation over the left FEF increased Multi-Scale Entropy in right frontal and bilateral parietal electrodes. Conversely, active *irregular* stimulation increased Multi-Scale Entropy over bilateral frontal and right parietal electrodes. Finally, in contrast with *non frequency-specific* patterns, 30 Hz *rhythmic* stimulation significantly increased Multi-Scale Entropy in miscellaneous electrodes located along the edge of the EEG grid, but failed to identify significant clusters of more than 3 electrodes.

These results suggest differences between 30 Hz *rhythmic* and the three *non frequency-specific* TMS patterns with regards to the modulation of Multi-Scale Entropy. However, direct two-by-two comparisons failed to show any significant differences in Multi-Scale Entropy between any of the four active TMS patterns probed in our study (Fig. 8C). See supplementary table S5 for full details on the statistical comparisons reported in Figures 8B and C.

### Impact on visual detection performance

As in prior studies (Chanes et al., 2013, 2015; Vernet et al., 2019), we explored the impact of 30 Hz *rhythmic* and the three *non frequency-specific* TMS conditions on conscious visual detection performance. To this end, we estimated potential changes of perceptual sensitivity (d’) across stimulation pattern types. We also explored shifts in response bias *via* decision criterion (c) and likelihood ratio (β) measures, both influenced by late perceptual decision-making processes.

Repeated-measures 2×2×4 ANOVAs with factors *Visual Field* (left, right), *TMS Condition* (active, sham) and *TMS Pattern* (*rhythmic, non-uniform rhythmic, random, irregular*) for perceptual sensitivity (d’) or likelihood ratio (β) did not yield any main effect or significant interaction (p>0.05). Our analyses, revealed however, a main effect of TMS condition on decision criterion (F(1,223)=9.154, MSE=0.136, p<0.05) suggesting that compared to sham TMS patterns, active TMS stimulation lowered decision criteria, hence made participants provide more liberal detection responses (i.e. be more likely to respond that a target was present on the screen in case of doubt). No main effect of *TMS Pattern*, nor interactions between *TMS Condition* and *TMS Pattern* was significant for decision criterion c (p>0.7), indicating that decision criterion remained unmodulated by the temporal distribution of the pulses present in any type of active TMS bursts.

## Discussion

We here aimed to investigate using non-invasive brain stimulation the modulation of internal neural noise in a left frontal region of the human brain, the left FEF, and explore potential top-down contribution of this cortical area to neurophysiological coding processes subtending the modulation of conscious visual perception. We did so by assessing the impact of episodic non frequency-specific TMS bursts on scalp EEG signals during a lateralized visual detection task for near-threshold stimuli. Our study extends to the neurophysiological domain prior behavioral evidence showing unexpected causal contributions of non frequency-specific left frontal activity to increases of visual sensitivity, which remained to be further validated and characterized by mean of combined TMS-EEG recordings (Chanes et al., 2015).

By calculating conventional EEG estimates of oscillation amplitude (power) and phase locking (ITC), we here showed that, surprisingly, none of the three *non frequency-specific* TMS patterns prevented a build-up of rhythmic cortical activity in the hight-beta range. In fact, rather paradoxically, the three stimulation patterns increased the power and aligned the phase of high-beta oscillations as much as 30 Hz rhythmic patterns (Thut et al., 2011b; Vernet et al., 2019). However, analyses also revealed that all three *non frequency-specific* patterns increased the power of left frontal oscillations across broader frequency bands, not necessarily restricted to ∼30 Hz. *Non uniform rhythmic* bursts induced a bimodal distribution of power in the high- and low-beta frequency bands, whereas, compared to rhythmic patterns, *irregular* bursts significantly increased phase locking within a wider band extending towards the low-beta range. Finally, a specific measure sensitive to levels of internal neural noise in EEG signals, Multi-Scale Entropy, suggested that the three types of *non frequency-specific* TMS patterns induced brain activity of higher complexity than *rhythmic* high-beta TMS bursts. Taken together, these outcomes suggest that *non frequency-specific* TMS patterns generate more heterogeneous broadband activity, and less predictable and more complex neural signals than a pure high-beta 30 Hz frequency.

Short bursts of arrhythmic or random TMS had been previously employed in multiple studies as control conditions to isolate the impact of frequency-specific components contained in rhythmic TMS bursts (Albouy et al., 2017; Chanes et al., 2013, 2015; Quentin et al., 2015; Stengel et al., 2021; Thut et al., 2011b; Vernet et al., 2019). However, their effect on ongoing electrophysiological signals and their potential cognitive contribution remained to be characterized in further detail by means of concurrent TMS-EEG recordings. Prior evidence has shown that the addition of optimal levels of peripheral noise (i.e. not too high but not too low levels of ‘dosed’ noise) to rhythmic visual stimuli could result in an enhancement of cortical oscillations entrained at this same frequency (Mori & Kai, 2002; Srebro & Malladi, 1999). Hence, plausibly, the three *non frequency-specific* patterns tested in our study could have acted similarly. Indeed, pulse onset timings emulated very closely a pure 30 Hz frequency. However, quite minor time shifts of the two middle pulses within a 4-pulse episodic TMS burst were likely responsible for adding noise to an underlying high-beta rhythmic activity.

The distribution of oscillation power across frequency bins from the power spectral density (PSD) analysis has been previously employed as a measure of entropy in electrophysiological signals (Rezek & Roberts, 1998). Frequencies distributed across a large number of frequency bins characterize unpredictable signals featuring higher entropy levels. This is in contrast with the synchronization of local oscillators at a single regular frequency during alpha or high-beta stimulation (Lin et al., 2021; Stengel et al., 2021; Thut et al., 2011a; Vernet et al., 2019) which gives rise to regular and predictable oscillating signals of very low entropy, within a single and narrow power peak. In this framework, EEG recordings in the present study showed that, compared to rhythmic bursts, the delivery of non frequency-specific TMS to the left frontal cortex resulted in higher levels of signal entropy, hence higher levels of neural noise. This conclusion was further strengthened by a second metric of neural noise, estimating changes of signal entropy over several time-scales, known as Multi-Scale Entropy (MSE). Higher values of Multi-Scale Entropy characterize a signal that is not random but complex and rich in information at several time-scales (Costa et al., 2002, 2005; Zhang, 1991). Accordingly, in our study, only *non frequency-specific* TMS patterns increased Multi-Scale Entropy over large clusters of scalp EEG contacts, compared to sham stimulation. Most interestingly, increases of signal complexity proved most significant for electrodes overlying right frontal and bilateral parietal regions, instead of those closest to the targeted left FEF site. The limited spatial resolution of scalp EEG precludes pinpointing the precise localization of the neural sources responsible for such effects on signal complexity. However, since the left FEF is an important node of a bilaterally distributed dorsal fronto-parietal network contributing to top-down attentional orienting (Corbetta et al., 2008), it is plausible that changes in signal complexity were enabled throughout this network as attentional processes occur during the time window in which TMS bursts were delivered, between the onset of the alerting cue and the appearance of the visual target (Gross et al., 2004; Kastner et al., 1999).

To the best of our knowledge, the outcomes of the current study offer first-time evidence that short TMS bursts locally modulate internal neural noise levels in a cortical site manipulated transcranially via focal neurostimulation. Furthermore, they provide experimental support in favor of the ability of TMS patterns with different types of temporal irregularities to induce distinct levels of internal cortical noise. Such effects could be considered similar to those hypothesized for transcranial Random Noise Stimulation (tRNS), a current-based stimulation technology able to mimick white noise signals (Terney et al., 2008), which for nearly a decade has been employed to explore and modulate different aspects of brain function. Transcranial RNS technology operates on the principle that by varying the amplitude of a randomly alternating current, different levels of neural noise can be cortically evoked (Groen et al., 2018; Groen & Wenderoth, 2016). In spite of some significant advantages compared to TMS (e.g., lower cost, ease of use, high portability and possibility of multi-site stimulation), tRNS suffers from the same limitations as other transcranial current stimulation (tCS) approaches; notably its low temporal and spatial resolution and a rather weak electrical field magnitude (Bikson et al., 2010; Datta et al., 2012; Nitsche et al., 2008). Additionally, concurrent tCS-EEG recordings remain very challenging and effective cleaning methods to remove electrical artifact generated by scalp-delivered electrical currents still rather controversial (Noury et al., 2016; Noury & Siegel, 2018). Consequently, compelling EEG evidence confirming the ability of tRNS to manipulate internal noise in ongoing neural signals is yet to be reported. In this scenario, the ability of *non frequency-specific* TMS bursts to operate focally and to induce time-circumscribed effects on EEG signals impacting specific behaviors holds particular promise to investigate coding strategies based on the generation and modulation of internal neural noise.

The Stochastic Resonance (SR) theory, presented decades ago, provides an insightful framework in which to further develop a mechanistic hypothesis for the TMS-EEG outcomes here reported. This body of evidence poses that the injection of optimal levels of noise in non-linear systems can boost the level of information carried by a signal (see Moss et al., 2004 for a review) and hence improve signal processing. Providing experimental support to this explanation, facilitatory effects of noise have been demonstrated at multiple levels of the nervous system. For example, the external addition of noise has shown to improve signal transduction through membrane ions channels (Bezrukov & Vodyanoy, 1995) and single cell responses to sensory stimuli (Collins et al., 1996; Cordo et al., 1996; Douglass et al., 1993; Jaramillo & Wiesenfeld, 1998). Moreover, in cortical regions, the episodic addition of stochastic noise to weak peripheral sensory stimuli increased evoked responses recorded by EEG (Srebro & Malladi, 1999) and enhanced cortical activity at the frequency in which such stimuli were delivered (Manjarrez et al., 2002; Mori & Kai, 2002). Lastly, at the behavioral level, the addition of noise to ongoing cortical activity facilitated the detection of weak sensory stimuli (Groen & Wenderoth, 2016; Iliopoulos et al., 2014; Kitajo et al., 2003; Manjarrez et al., 2007) and led to improvements in coding of higher cognitive processes such as decision-making or memory (Groen et al., 2018; Usher & Feingold, 2000).

Although our TMS-EEG measures showed an impact of stimulation on internal noise levels, we failed to replicate prior improvements of visual sensitivity driven by two of our three *non frequency-specific* TMS bursts (*non-uniform rhythmic* and *random*) delivered to the left FEF (Chanes et al., 2015). Several reasons could account for this outcome. First, detection improvements for visual stimuli with the addition of stochastic noise have been shown to follow an inverse U-shaped curve (Collins et al., 1996; Simonotto et al., 1997). Accordingly, the addition of proper levels of noise drives detection improvements, whereas quantities slightly below or above have either no effect or a detrimental impact on neural signals, and hence may fail to significantly modulate behavioral performance. Most importantly, prior literature strongly suggests that the optimal levels of external noise to drive behavioral improvements vary substantially across individuals (Iliopoulos et al., 2014; Kitajo et al., 2003), and that the presence of high levels of ongoing internal noise may weaken facilitatory phenomena tied to Stochastic Resonance (Aihara et al., 2008). On this basis, we cannot rule out the possibility that fixed patterns of noise carried by some of the tested *non frequency-specific* TMS patterns (*non-uniform rhythmic*, *irregular* and *random*) might have been either too low or too high to improve visual perception in individual participants, due to differences in their individual level of ongoing internal noise. Moreover, even if noise was induced in a range susceptible to engage behavioral improvement, inter-individual variability in the required levels of optimal noise could have extended the facilitatory effect to some or all of our three *non frequency-specific* TMS patterns, cancelling off statistical differences between them at the group level (Groen & Wenderoth, 2016).

In any case, the lack of significant pattern-specific perceptual outcomes must encourage a search for approaches and metrics guiding the customization of external noise levels able to boost internal neural signals, improve cortical processing and ultimately enhance visual performance. To this regard, the use of longer TMS bursts made of more than four pulses or, when behaviorally relevant, delivered at slower TMS frequencies could provide additional freedom to tailor the structure of a burst across a *continuum* between complete pulse onset randomness and pure and perfectly regular rhythmic oscillations. Higher flexibility to titrate noise levels would allow individualization of non frequency-specific TMS patterns, leading to lower inter-subject variability, and a more robust impact on visual performance at the group level.

Overall, in this study, we probed hypotheses concerning the impact of non frequency-specific TMS bursts on brain activity assessed with EEG signals (Chanes et al., 2015). We brought evidence that such patterns increased the bandwidth of local neural oscillations and generated signals of lower predictability and higher complexity in the targeted left frontal regions (FEF) and interconnected left and right fronto-parietal cortical sites. Together with previously published studies demonstrating a causal impact of short TMS patterns on visual detection performance (Chanes et al., 2013, 2015; Stengel et al., 2021; Vernet et al., 2019), the current results also support an asymmetry of coding strategies between the left and right dorsal attentional systems for the top-down modulation of conscious visual perception; with evidence of a causal role of high-beta oscillations in right fronto-parietal systems (Chanes et al., 2013; Stengel et al., 2021; Vernet et al., 2019) and the contribution of ‘dosed’ levels of neural noise (or non-predictable activity patterns) in left hemisphere homotopic networks (Chanes et al., 2015, current results). This anatomical and functional model would need to be further validated ideally with an *ad hoc* experiment assessing frequency-specific and non frequency-specific TMS patterns on both the left and right FEF in the same population of participants, hence remains at this point speculative.

Importantly, we should not fall into the pitfall of considering oscillations and neural noise as two segregated classes of opposite − or mutually excluding − brain activity patterns involved in cognitive coding. Early evidence supporting Stochastic Resonance theory reported counter-intuitively that the addition of noise could result in unexpected increases of highly regular and predictable activity (Mori & Kai, 2002; Srebro & Malladi, 1999). Hence on the basis of this framework and considering the current data, we here aim to conceptualize both oscillations and neural noise as two neurophysiological strategies representing the ends of a long continuum contributing to common modulatory mechanisms of brain activity. Accordingly, in physiological conditions, both the entrainment of oscillations or the increase of internal neural noise may jointly contribute to enhance or suppress cerebral coding strategies enabling specific cognitive or behavioral events. Likewise, during non-invasive brain stimulation, external bursts of rhythmic or non frequency-specific stimulation interact with ongoing patterns of internal neural noise levels within the targeted cortical region. Consequently, net modulatory effects depend on the balance between internal and externally added sources of noise. Since cortical neural noise levels have been shown to vary greatly between individuals, those with naturally higher cortical noise levels are less likely to show stochastic resonance-like improvements of stimulus detection with the addition of external noise (Aihara et al., 2008). We here speculate that similar effects might be at play in left and right dorsal fronto-parietal attention networks, which we manipulated in this and prior studies, respectively (Chanes et al., 2013, 2015; Stengel et al., 2021; Vernet et al., 2019). In neural networks with a higher level of ongoing neural noise level, the entrainment of regular cortical oscillations (i.e., signals with scarce noise) would most contribute to improvements of visual detection. However, in the opposite scenario, i.e., sites and networks with very low levels of ongoing neural noise, it is the addition of non-predictable activity that would be more prone to induce stochastic resonance-like improvements of signal detection and be most likely to affect behavioral outputs. Doubtless, the identification of reliable measures of ongoing internal neural noise levels throughout cortical areas which could be used to dose external noise levels via brain stimulation will provide a basis to further validate this hypothetical physiological model.

In conclusion, using non-invasive causal approaches we here demonstrated the ability of non frequency-specific brain stimulation bursts to modulate levels of neural noise, measured in terms of frequency heterogeneity (Frequency Peak Width), predictability (Sample Entropy) and signal complexity (Multiscale Sample Entropy). Yet, at difference with prior observations (Chanes et al., 2015), we were unable to show significant pattern-specific modulations of visual performance outcomes. This is likely due to inter-individual differences in internal noise levels, an outcome emphasizing the need to customize interventions according to ongoing internal noise levels. On the basis of our findings and their present and future implications, our study encourages further work in at least three different directions: first, to better characterize the role of neural noise in interaction with oscillatory activity in the processing of cortical signals and the coding of specific cognitive events, identifying the sources and mechanisms of internal neural noise; second, to further explore the ability of well-dosed transcranially-induced neural noise to facilitate visual perception outcomes by boosting attentional orienting networks, and further understand its underpinning mechanisms; third and last, to develop reliable estimates and online monitoring of ongoing levels of internal noise in order to enable the customization of non frequency-specific brain stimulation patterns, accounting for individual variability.

## Acknowledgements

Chloé Stengel was supported by a PhD fellowship from the University Pierre and Marie Curie. Julià L. Amengual was supported by a fellowship from the *Fondation Fyssen*. The activities of the laboratory of Dr. Valero-Cabré are supported by research grants IHU-A-ICM-Translationnel, ANR projet Générique OSCILOSCOPUS and Flag-Era-JTC-2017 CAUSALTOMICS. The authors would also like to thank Marine Vernet and Clara Sanches for their help during initial and late phases of data acquisition and the Naturalia & Biologia Foundation for financial assistance for traveling and attending meetings.

## Author contributions

Conceptualization: A.V-C and C.S. Data acquisition: C.S. and A.V.-C. Data analysis and data interpretation: C.S. and A.V-C. Manuscript preparation: C.S, A.V-C, J.L.A & T.M. General supervision: A.V-C.

## Data availability statement

Data are available from the corresponding author upon request.

