## Supplementary Materials for "Causal contributions of neural noise to left frontal networks subtending the modulation of conscious visual perception in the human brain"

Chloé Stengel[1]*, Julià L. Amengual[1,2], Tristan Moreau[1] & Antoni Valero-Cabré [1,3,4]*

[1] Cerebral Dynamics, Plasticity and Rehabilitation Group, FRONTLAB, Institut du Cerveau, CNRS UMR 7225, Paris, France

[2] Institut des Sciences Cognitives Marc Jeannerod, Département de Neuroscience Cognitive, CNRS UMR 5229, Université Claude Bernard Lyon I, 67 Boulevard Pinel, 69675 Bron Cedex, France.

[3] Dept. Anatomy and Neurobiology, Laboratory of Cerebral Dynamics, Boston University School of Medicine, Boston, USA

[4] Cognitive Neuroscience and Information Tech. Research Program, Open University of Catalonia (UOC), Barcelona, SPAIN

[*] Corresponding authors: Antoni Valero-Cabré, MD PhD & Chloé Stengel PhD

Groupe de Dynamiques Cérébrales, Plasticité et Rééducation

Équipe FRONTLAB, Institut du Cerveau (ICM)

Hôpital de la Pitié-Salpêtrière, 47 boulevard de l'Hôpital, 75013 Paris, France.

 &

**Supplementary analyses**

Similarly to what was presented for the analysis of power-spectrum peaks in the main article (Fig. 7), we investigated peak-width of increased phase-alignment during active stimulation. This analysis followed the same method described for power-spectrum peaks, namely, local maxima were identified on ITC profiles computed over frequency through a jackknife procedure during the whole stimulation window ([-133 0]ms, time centered on visual target onset). The peak frequency and peak-width of each local maxima was then extracted.

Comparison of ITC peak-width between active TMS patterns produced similar results as those reported for power peak-width in the main article. The *non-uniform rhythmic* active TMS pattern differed from the 3 other patterns in that 2 phase-alignment peaks could be identified on the ITC profile (Fig. S1A). One high-beta peak at ~35 Hz was identified in all 15 iterations of the jackknife procedure and one low-beta peak at ~13 Hz was identified in 13 of the 15 iterations of the jackknife procedure. For all other active TMS patterns, a single peak was identified in the high-beta range: at ~30 Hz for the active *rhythmic* condition, at ~28 Hz for the active *random* condition and at ~25 Hz for the active *irregular* condition.

The frequency bandwith of the increases in oscillatory phase-alignment for the three TMS patterns showing a single high-beta peak was compared using a one-way ANOVA. A significant difference across active *rhythmic*, *random* and *irregular* stimulation patterns was found ($F(2,41) = 4.234$, $MSE=0.6$, $p<0.05$, after correction for reduced error variance from the jackknife procedure). Post-hoc paired Student *t*-test showed that active *irregular* TMS increased oscillation phase-alignment in a significantly broader frequency band than *rhythmic* ($T(14)=5.788$, $p<0.01$, corrected for reduced error variance by jackknife procedure) or *random* TMS bursts ($T(14) = 2.108$, $p<0.05$, corrected for reduced error variance by jackknife procedure) (Fig S1B).

Overall, this analysis confirms the results of the analysis on power peak-width presented in the main article by showing that *non frequency-specific* TMS patterns modulated oscillatory activity in wider and more diverse frequency bands than the *rhythmic* pattern.

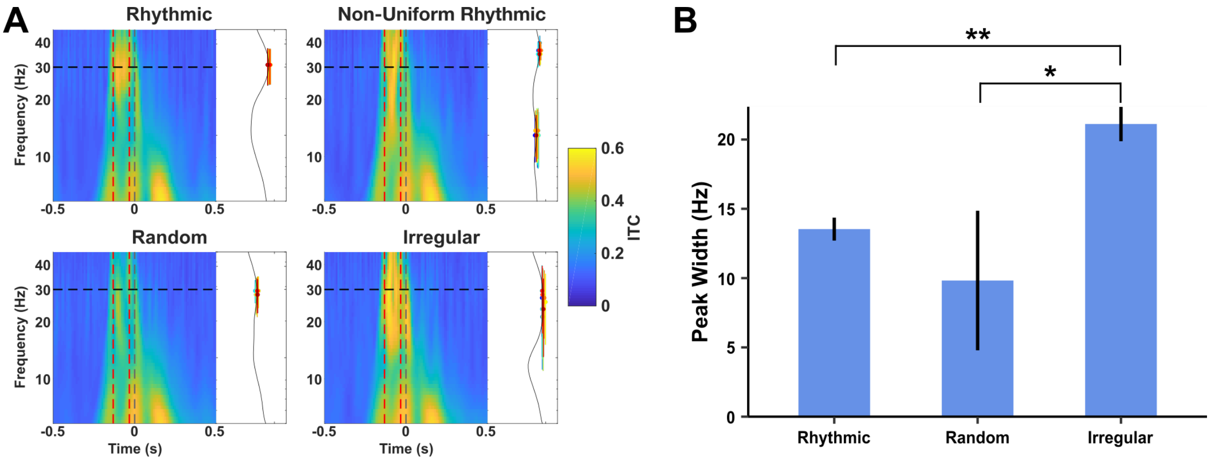


**Supplementary Figure S1. Peak-width of phase-alignment increases during 30 Hz rhythmic TMS compared to the three non frequency-specific TMS patterns. (A)** Time-frequency maps in a cluster of left frontal electrodes (F1, F3, FC1, FC3) closest to the center of the stimulation coil during active stimulation trials represented for each stimulation pattern (30 Hz *rhythmic* TMS, and three *non frequency-specific* patterns: *non-uniform rhythmic*, *random* and *irregular*). The two red dotted vertical lines signal the first (-133 ms) and last (-33 ms) TMS pulses of the burst. The black dotted horizontal line indicates the frequency (30 Hz) of the rhythmic TMS pattern. Right marginal graphs of each time-frequency panel display the average ITC spectrum over the whole window of TMS delivery. Colored lines show the width of the peaks of ITC as detected with a jackknife procedure. **(B)** Comparison of width of ITC peaks (expressed in Hz) during active stimulation between *rhythmic* and *non frequency specific* (*random* and *irregular*) TMS. The so called *non-uniform rhythmic* TMS pattern is not presented in the figure, because its ITC spectrum during active TMS revealed two distinct peaks (35 Hz and 13 Hz, panel A) and thus could not be directly compared to those displaying only a single peak in the high-beta range (~30 Hz). Notice that error bars represent the standard error corrected for the reduced error variance by the jackknife procedure. One-way ANOVA analysis yielded a significant main effect of TMS pattern. Notice that *irregular* TMS patterns increased cortical oscillatory phase-alignment in a significantly wider frequency band than *rhythmic* or *random* TMS patterns. Results of the post-hoc t-tests are indicated as follows: ** $p < 0.01$, * $p < 0.05$.

**Supplementary Tables**

**Supplementary Table S1: Statistical results for cluster-based permutation tests on high-beta power topography during stimulation.** Corresponding to statistical results shown in Figure 3 of the main article text.

| Comparison | Cluster number | Electrodes in cluster | Cluster size (nb of electrodes) | Cluster T-statistic | Cluster p-value |
|---|---|---|---|---|---|
| **Rhythmic Active vs. Sham** | 1 | FP1, FPz, AF7, AF3, AFz, AF4, F7, F5, F3, F1, Fz, F2, F4, F6, FT7, FC5, FC3, FC1, FCz, FC2, FC4, FC6, FT8, C5, C3, C1, Cz, C2, C4, C6, TP7, CP5, CP3, CP1, CPz, CP2, CP4, CP6, TP8, P7, P5, P3, P1, Pz, P2, P4, P6, P8, PO7, PO3, POz, PO4, PO8, O1, Oz, O2, Iz | 57 | 260.743142 | 0.000100 |
| **Non-uniform rhythmic Active vs. Sham** | 1 | FP1, FPz, FP2, AF7, AF3, AFz, AF4, F7, F5, F3, F1, Fz, F2, F4, F6, F8, FT7, FC5, FC3, FC1, FCz, FC2, FC4, FC6, FT8, C5, C3, C1, Cz, C2, C4, C6, TP7, CP5, CP3, CP1, CPz, CP2, CP4, CP6, TP8, P7, P5, P3, P1, Pz, P2, P4, P6, P8, PO7, PO3, POz, PO4, PO8, O1, Oz, O2, Iz | 59 | 287.402805 | 0.000100 |
| **Random Active vs. Sham** | 1 | FP1, FPz, FP2, AF7, AF3, AFz, AF4, AF8, F7, F5, F3, F1, Fz, F2, F4, F8, FT7, FC5, FC3, FC1, FCz, FC2, FC4, FC6, FT8, C5, C3, C1, Cz, C2, C4, C6, TP7, CP5, CP3, CP1, CPz, CP2, CP4, CP6, TP8, P7, P5, P3, P1, Pz, P2, P4, P6, P8, PO7, PO3, POz, PO4, PO8, O1, Oz, O2, Iz | 59 | 288.810225 | 0.000100 |
| **Irregular Active vs. Sham** | 1 | FP1, FPz, FP2, AF7, AF3, AFz, AF4, AF8, F7, F5, F3, F1, Fz, F2, F4, F6, F8, FT7, FC5, FC3, FC1, FCz, FC2, FC4, FC6, FT8, C5, C3, C1, Cz, C2, C4, C6, TP7, CP5, CP3, CP1, CPz, CP2, CP4, CP6, TP8, P7, P5, P3, P1, Pz, P2, P4, P6, P8, PO7, PO3, POz, PO4, PO8, O1, Oz, O2, Iz | 60 | 299.816992 | 0.000100 |

Note that none of the direct statistical comparisons between active TMS patterns yielded any significant clusters (no electrodes survived the alpha = 0.01 threshold for student's t-test) therefore no cluster-based permutation tests could be performed for these comparisons.

**Supplementary Table S2: Statistical results for cluster-based permutation tests on high-beta ITC topography.** Corresponding to statistical results shown in Figure 4 of the main article text.

| Comparison | Cluster number | Electrodes in cluster | Cluster size (nb of electrodes) | Cluster T-statistic | Cluster p-value |
|---|---|---|---|---|---|
| **Rhythmic Active vs. Sham** | 1 | FP1, FPz, FP2, AF7, AF3, AFz, AF4, AF8, F7, F5, F3, F1, Fz, F2, F4, F6, F8, FT7, FC5, FC3, FC1, FCz, FC2, FC4, FC6, FT8, C5, C3, C1, Cz, C2, C4, C6, TP7, CP5, CP3, CP1, CPz, CP2, CP4, CP6, TP8, P7, P5, P3, P1, Pz, P2, P4, P6, P8, PO7, PO3, POz, PO4, PO8, O1, Oz, O2, Iz | 60 | 322.846036 | 0.000100 |
| **Non-uniform rhythmic Active vs. Sham** | 1 | FP1, FPz, FP2, AF7, AF3, AFz, AF4, AF8, F7, F5, F3, F1, Fz, F2, F4, F6, F8, FT7, FC5, FC3, FC1, FCz, FC2, FC4, FC6, FT8, C5, C3, C1, Cz, C2, C4, C6, TP7, CP5, CP3, CP1, CPz, CP2, CP4, CP6, TP8, P7, P5, P3, P1, Pz, P2, P4, P6, P8, PO7, PO3, POz, PO4, PO8, O1, Oz, O2, Iz | 60 | 429.680676 | 0.000100 |
| **Random Active vs. Sham** | 1 | FP1, FPz, FP2, AF7, AF3, AFz, AF4, AF8, F7, F5, F3, F1, Fz, F2, F4, F6, F8, FT7, FC5, FC3, FC1, FCz, FC2, FC4, FC6, FT8, C5, C3, C1, Cz, C2, C4, C6, TP7, CP5, CP3, CP1, CPz, CP2, CP4, CP6, TP8, P7, P5, P3, P1, Pz, P2, P4, P6, P8, PO7, PO3, POz, PO4, PO8, O1, Oz, O2, Iz | 60 | 334.524987 | 0.000100 |
| **Irregular Active vs. Sham** | 1 | FP1, FPz, FP2, AF7, AF3, AFz, AF4, AF8, F7, F5, F3, F1, Fz, F2, F4, F6, F8, FT7, FC5, FC3, FC1, FCz, FC2, FC4, FC6, FT8, C5, C3, C1, Cz, C2, C4, C6, TP7, CP5, CP3, CP1, CPz, CP2, CP4, CP6, TP8, P7, P5, P3, P1, Pz, P2, P4, P6, P8, PO7, PO3, POz, PO4, PO8, O1, Oz, O2, Iz | 60 | 429.694351 | 0.000100 |
| **Active Rhythmic vs. Random** | 1 | F5, F3, FT7, FC5, FC3, FC1, C5, C3, TP7, CP5, CP3, P7, P5, P3, P1, Pz, PO7, PO3, POz, O1 | 20 | 74.363274 | 0.000300 |
| **Active Rhythmic vs. Irregular** | 1 | F2, FCz | 2 | -7.358359 | 0.016998 |
| | 2 | O2 | 1 | -3.132227 | 0.058394 |
| **Active Non-uniform rhythmic vs. Random** | 1 | FP1, AF3, AFz, F7, F5, F3, F1, Fz, F2, F4, F6, FT7, FC3, FCz, FC2, FC4, FC6, FT8, C3, Cz, C2, C4, C6, CP5, CP3, CPz, CP2, CP4, CP6, P7, P5, P3, P1, Pz, P2, P4, P6, P8, PO7, PO3, POz, PO4, PO8 | 43 | 158.940738 | 0.000900 |
| **Active Non-uniform rhythmic vs. Irregular** | 1 | C5 | 1 | -3.177122 | 0.062394 |
| | 2 | FC3 | 1 | -3.169035 | 0.062994 |
| **Active Random vs. Irregular** | 1 | FP1, FPz, FP2, AF7, AF3, AFz, AF4, F7, F5, F3, F1, Fz, F2, F4, F6, F8, FT7, FC5, FC3, FC1, FCz, FC2, FC4, FC6, C5, C3, C1, Cz, C2, C4, C6, TP7, CP5, CP3, CP1, CPz, CP2, CP4, CP6, TP8, P7, P5, P3, P1, Pz, P2, P4, P6, P8, PO7, PO3, POz, PO4, PO8, O1, Oz, O2, Iz | 58 | -282.21843 | 0.000100 |

**Supplementary Table S3: Statistical results for cluster-based permutation tests on left frontal power time-frequency analysis.** Corresponding to statistical results shown in Figure 5 of the main article text.

| Comparison | Cluster number | Time (s) [min max] | Frequency (Hz) [min max] | Cluster size (nb of time-frequency points) | Cluster T-statistic | Cluster p-value |
|---|---|---|---|---|---|---|
| **Rhythmic Active vs. Sham** | 1 | [-0.13 0.1] | [11 45.75] | 484 | 2120.221082 | 0.001000 |
| | 2 | [-0.18 -0.08] | [6 9.75] | 92 | 352.694407 | 0.016298 |
| | 3 | [0.12 0.18] | [6 8] | 37 | 118.976757 | 0.064694 |
| | 4 | [-0.27 -0.23] | [13.75 21] | 31 | 111.631006 | 0.069793 |
| **Non-uniform rhythmic Active vs. Sham** | 1 | [-0.19 0.2] | [6 45.75] | 952 | 4019.306721 | 0.000400 |
| **Random Active vs. Sham** | 1 | [-0.17 0.2] | [6 45.75] | 929 | 4139.041934 | 0.000100 |
| | 2 | [-0.3 -0.26] | [9.25 12.5] | 34 | -115.906010 | 0.064294 |
| | 3 | [-0.23 -0.22] | [16.25 20.25] | 10 | -30.706000 | 0.187581 |
| **Irregular Active vs. Sham** | 1 | [-0.3 0.2] | [6 45.75] | 1037 | 4090.949675 | 0.000100 |
| **Active rhythmic vs. Non-uniform rhythmic** | 1 | [-0.14 -0.1] | [9.75 13] | 30 | -97.986708 | 0.077492 |
| | 2 | [-0.26 -0.25] | [13.75 16.25] | 8 | 25.128227 | 0.270273 |
| **Active Rhythmic vs. Random** | 1 | [-0.27 -0.21] | [13 24] | 74 | 265.705441 | 0.019098 |
| | 2 | [0.15 0.18] | [19.25 29.75] | 28 | 97.174479 | 0.092991 |
| **Active Rhythmic vs. Irregular** | 1 | [-0.25 -0.23] | [18.5 33.75] | 38 | 165.011315 | 0.041096 |
| | 2 | [-0.09 -0.05] | [14.25 19.25] | 32 | -107.679371 | 0.064594 |
| | 3 | [-0.08 -0.06] | [33.75 45.75] | 18 | -64.305259 | 0.100290 |
| **Active Non-uniform rhythmic vs. Random** | 1 | [-0.14 -0.12] | [33.75 42] | 13 | 42.521708 | 0.212079 |
| | 2 | [-0.22 -0.21] | [17.75 23] | 12 | 38.599607 | 0.226077 |
| | 3 | [-0.02 -0.01] | [32.5 45.75] | 11 | 38.448337 | 0.226977 |
| | 4 | [-0.04 -0.03] | [20.25 25] | 11 | -37.099955 | 0.240676 |
| | 5 | [-0.09 -0.08] | [24 31] | 10 | -35.616362 | 0.244876 |
| **Active Non-uniform rhythmic vs. Irregular** | 1 | [-0.16 -0.11] | [7.75 16.25] | 81 | 323.910749 | 0.010699 |
| | 2 | [-0.11 -0.06] | [16.25 35.25] | 56 | -218.737313 | 0.024998 |
| | 3 | [-0.19 -0.17] | [26.25 35.25] | 11 | 36.462468 | 0.220278 |
| | 4 | [-0.01 -0.01] | [37 45.75] | 6 | 18.919307 | 0.329167 |
| | 5 | [-0.12 -0.12] | [32.5 37] | 4 | 14.253149 | 0.372463 |
| **Active Random vs. Irregular** | 1 | [-0.09 -0.06] | [13.75 18.5] | 21 | -70.113892 | 0.136586 |
| | 2 | [-0.08 -0.06] | [37 45.75] | 13 | -43.687156 | 0.191881 |

**Supplementary Table S4: Statistical results for cluster-based permutation tests on left frontal ITC time-frequency analysis.** Corresponding to statistical results shown in Figure 6 of the main article text.

| Comparison | Cluster number | Time (s) [min max] | Frequency (Hz) [min max] | Cluster size (nb of time-frequency points) | Cluster T-statistic | Cluster p-value |
|---|---|---|---|---|---|---|
| **Rhythmic Active vs. Sham** | 1 | [-0.21 0.2] | [6 45.75] | 983 | 5150.862711 | 0.000100 |
| | 2 | [0.16 0.16] | [20.25 40.25] | 17 | 61.871736 | 0.115088 |
| | 3 | [-0.21 -0.2] | [27.25 37] | 10 | 35.430008 | 0.187881 |
| | 4 | [0.13 0.13] | [28.5 38.5] | 8 | 26.653448 | 0.233177 |
| **Non-uniform rhythmic Active vs. Sham** | 1 | [-0.22 0.2] | [6 45.75] | 1093 | 5483.605946 | 0.000100 |
| | 2 | [0.1 0.1] | [26.25 35.25] | 8 | 26.973565 | 0.238276 |
| | 3 | [-0.27 -0.27] | [24 32.5] | 8 | 25.632726 | 0.248375 |
| **Random Active vs. Sham** | 1 | [-0.21 0.2] | [6 45.75] | 1135 | 5732.008936 | 0.000100 |
| | 2 | [-0.21 -0.2] | [18.5 31] | 20 | 82.035202 | 0.078592 |
| | 3 | [0.07 0.07] | [29.75 45.75] | 11 | 47.650280 | 0.133387 |
| | 4 | [-0.17 -0.17] | [26.25 33.75] | 7 | 24.619939 | 0.229077 |
| | 5 | [0.17 0.17] | [28.5 37] | 7 | 23.929423 | 0.232677 |
| **Irregular Active vs. Sham** | 1 | [-0.21 0.2] | [6 45.75] | 1075 | 5483.103530 | 0.000100 |
| | 2 | [-0.3 -0.3] | [19.25 28.5] | 10 | 36.054534 | 0.180782 |
| | 3 | [-0.27 -0.27] | [31 40.25] | 7 | 24.033447 | 0.242376 |
| | 4 | [-0.25 -0.25] | [22 28.5] | 7 | 22.351621 | 0.251975 |
| **Active rhythmic vs. Non-uniform rhythmic** | 1 | [-0.14 -0.1] | [11 16.25] | 35 | -141.602107 | 0.034897 |
| | 2 | [0.07 0.09] | [6 8.5] | 16 | -51.150808 | 0.180682 |
| **Active Rhythmic vs. Random** | 1 | [-0.07 -0.04] | [21 45.75] | 62 | 264.069147 | 0.007999 |
| | 2 | [0.03 0.09] | [6 6.5] | 16 | -54.241227 | 0.179482 |
| **Active Rhythmic vs. Irregular** | 1 | [-0.12 -0.03] | [12.5 23] | 102 | -368.148203 | 0.002500 |
| | 2 | [-0.01 0.02] | [15.5 27.25] | 35 | -132.441098 | 0.046395 |
| **Active Non-uniform rhythmic vs. Random** | 1 | [-0.12 -0.04] | [22 45.75] | 75 | 318.336090 | 0.006099 |
| | 2 | [-0.13 -0.11] | [11 16.25] | 23 | 83.003023 | 0.088391 |
| | 3 | [0.13 0.13] | [28.5 32.5] | 4 | 13.282001 | 0.503550 |
| **Active Non-uniform rhythmic vs. Irregular** | 1 | [-0.14 -0.1] | [19.25 29.75] | 29 | -126.518532 | 0.046595 |
| | 2 | [0.16 0.18] | [9.25 16.25] | 26 | 89.082071 | 0.084892 |
| | 3 | [-0.15 -0.12] | [9.75 13.75] | 19 | 63.602808 | 0.142686 |
| | 4 | [0 0.01] | [18.5 24] | 12 | -46.451610 | 0.210179 |
| | 5 | [0.13 0.13] | [29.75 38.5] | 7 | 25.202883 | 0.375162 |
| | 6 | [-0.04 -0.04] | [17.75 20.25] | 4 | -12.818915 | 0.533547 |
| **Active Random vs. Irregular** | 1 | [-0.14 0.03] | [14.25 45.75] | 285 | -1225.581050 | 0.000100 |
| | 2 | [-0.24 -0.23] | [8.5 9.25] | 5 | -15.633403 | 0.455554 |

**Supplementary Table S5: Statistical results for cluster-based permutation tests on Multi-Scale Entropy (MSE) topography during stimulation.** Corresponding to statistical results shown in Figure 8, panel B and C, of the main article text.

| Comparison | Cluster number | Electrodes in cluster | Cluster size (nb of electrodes) | Cluster T-statistic | Cluster p-value |
|---|---|---|---|---|---|
| **Rhythmic Active vs. Sham** | 1 | FT7, C5, CP5 | 3 | 9.353347 | 0.015598 |
| | 2 | C6, TP8 | 2 | 6.797519 | 0.016898 |
| | 3 | AF8, F8 | 2 | 6.178491 | 0.019298 |
| | 4 | Oz | 1 | 3.001531 | 0.038996 |
| **Non-uniform rhythmic Active vs. Sham** | 1 | AF4, Fz, F2, F4, FC2, FC4, FC6, C2, C4, C6, CP2, CP4, CP6, TP8, P4 | 15 | 49.478339 | 0.002500 |
| | 2 | F7, FT7, C5, CP5, CP3, P7, P5, P3, P1, Pz, PO7, PO3, POz, Oz | 14 | 45.918899 | 0.003200 |
| | 3 | FC1 | 1 | 3.310926 | 0.026297 |
| **Random Active vs. Sham** | 1 | AF4, Fz, F2, F4, F6, F8, FC4, FC6, C6, TP8 | 10 | 35.650535 | 0.006399 |
| | 2 | FT7, C5, CP5, P5 | 4 | 14.133527 | 0.010199 |
| **Irregular Active vs. Sham** | 1 | FP2, AF4, AF8, F7, F5, F3, F1, Fz, F2, F4, F6, F8, FT7, FC1, FCz, FC2, FC4, FC6, FT8, C5, C2, C4, C6, CP5, CP2, CP4, CP6, TP8, Pz, P2, P4, P6, P8, PO3, POz, PO4, Oz, O2, Iz | 39 | 145.876917 | 0.000900 |
| **Active Rhythmic vs. Non-uniform rhythmic** | 1 | AF8 | 1 | 3.155999 | 0.097290 |
| **Active Rhythmic vs. Irregular** | 1 | FT8 | 1 | -3.209091 | 0.073593 |
| **Active Non-uniform rhythmic vs. Random** | 1 | C5 | 1 | -2.989813 | 0.085191 |
| **Active Non-uniform rhythmic vs. Irregular** | 1 | AF8 | 1 | -3.246434 | 0.054695 |
